# Structural and biochemical characterization of LIG1 during mutagenic nick sealing of oxidatively damaged ends at the final step of DNA repair

**DOI:** 10.1101/2024.05.06.592774

**Authors:** Kanal Elamparithi Balu, Danah Almohdar, Jacob Ratcliffe, Qun Tang, Tanay Parwal, Melike Çağlayan

## Abstract

DNA ligase 1 (LIG1) joins broken strand-breaks in the phosphodiester backbone to finalize DNA repair pathways. We previously reported that LIG1 fails on nick repair intermediate with 3’-oxidative damage incorporated by DNA polymerase (pol) β at the downstream steps of base excision repair (BER) pathway. Here, we determined X-ray structures of LIG1/nick DNA complexes containing 3’-8oxodG and 3’-8oxorG opposite either a templating Cytosine or Adenine and demonstrated that the ligase active site engages with mutagenic repair intermediates during steps 2 and 3 of the ligation reaction referring to the formation of DNA-AMP intermediate and a final phosphodiester bond, respectively. Furthermore, we showed the mutagenic nick sealing of DNA substrates with 3’-8oxodG:A and 3’-8oxorG:A by LIG1 wild-type, immunodeficiency disease-associated variants, and DNA ligase 3α (LIG3α) *in vitro*. Finally, we observed that LIG1 and LIG3α seal resulting nick after an incorporation of 8oxorGTP:A by polβ and AP-Endonuclease 1 (APE1) can clean oxidatively damaged ends at the final steps. Overall, our findings uncover a mechanistic insight into how LIG1 discriminates DNA or DNA/RNA junctions including oxidative damage and a functional coordination between the downstream enzymes, polβ, APE1, and BER ligases, to process mutagenic repair intermediates to maintain repair efficiency.

## Introduction

Genomic DNA can be oxidized by the reactive oxygen species (ROS) generated as byproducts of oxygen respiration or molecular executors in the host defense, and by various endogenous stressors and exposure to environmental hazards such as ionizing radiation and chemical toxicants (1–3). DNA precursor nucleotides (dNTPs) in the cellular nucleotide pool are subject to oxidative damage during oxidative stress, inflammation, and environmental exposure (4–6). Mammalian cells sanitize oxidized nucleotides by hydrolyzing them into monophosphate forms through the enzymatic activity of MutT homolog 1 (MTH1), and tumors have higher levels of MTH1 than adjacent healthy tissues and the chemical inhibition of MTH1 has been widely used to target cancer cells to elevate 8oxodG incorporation into genome (7–10). Guanine is particularly more susceptible to oxidation and 8-oxo-2’-deoxyguanine (8-oxodG) is the most abundant and the best characterized oxidized base as a biomarker of DNA oxidation (11–13). 8oxodG forms base pairing with Cytosine in *anti*-conformation and adopts *syn*-conformation when pairing with Adenine (14). Similarly, the most frequently oxidized nucleotide is dGTP in DNA precursor pool, and 7,8-dihydro-8’-oxo-dGTP (8oxodGTP) is also incorporated into the template strand opposite either Cytosine (nonmutagenic) or Adenine (mutagenic) and the latter is implicated in driving tumorigenesis causing A-T to C-G transversions (15–20). Ribonucleotides (rNTPs), the most abundant non-canonical nucleotides, can be frequently incorporated into genome at a remarkably high rate, impacting genome stability (21). Oxidation of cellular rNTP pools by ROS generates oxidized ribonucleotides and rGTP is analogously oxidized to 7,8-dihydro-8-oxo-guanosine (8-oxorGTP) (22,23). Oxidized nucleotides can interfere with faithful DNA replication or transcription leading to genome instability and induce various cellular abnormalities, such as mutations, cancer, neurological diseases, and cellular senescence (3,5,6,10,12,14,15). During times of oxidative stress, repair and replication DNA polymerases from A, B, C, X, and Y families, can perform futile DNA synthesis by inserting 8oxodGTP and 8oxorGTP, preferentially opposite a favorable template base Adenine, with different catalytic efficiency and fidelity (24–41).

Playing essential roles in nuclear replication, DNA repair, and recombination, human DNA ligases, DNA ligase 1 (LIG1), DNA ligase 3α (LIG3α), and DNA ligase IV (LIGIV), catalyze a metal-dependent phosphodiester bond formation between adjacent 3’-hydroxyl (3’-OH) and 5’-phosphoryl termini (5’-PO_4_) in duplex DNA via a similar mechanism and the ligation fidelity prerequisites the presence of proper DNA ends for faithful nick sealing (42–44). Incorporation of mismatches or oxidized nucleotides by error-prone DNA polymerases during DNA synthesis step of DNA repair would give rise to formation of nicks containing 3’-mismatched or oxidatively damaged ends at the final step (45–47). DNA ligases catalyze a highly conserved ligation reaction involving three sequential step by utilizing the energy of ATP (48): the formation of the ligase:adenylate intermediate after the active-site lysine residue (K568 of LIG1) attacks and forms a covalent bond with the α-phosphate of ATP in step 1. AMP is then transferred from the adenylated ligase to the 5’-PO_4_ end of the nick DNA, generating a 5’-adenylated DNA intermediate during step 2. Finally, the 3’-OH terminus attacks the 5’-P terminus downstream of the nick so that DNA ligase catalyzes a phosphodiester bond formation between adjacent ends, which is coupled to release of AMP in step 3 of the ligation reaction.

LIG1, the main replicative ligase, completes maturation of the Okazaki fragments by ligating >50 million nicks during DNA replication and finalizes DNA excision repair pathways by an ultimate ligation step (49). It has been shown that LIG1 discriminates against DNA junctions harboring mismatches and oxidative DNA damage at the 3’-end of nick and employs Mg^2+^-reinforced DNA binding to ensure high-fidelity (50,51). Our structures further uncovered the strategies that the ligase active site adopts to deter or favor a mis-paired terminus:template such as G:T and A:C mismatches (52). We recently demonstrated that LIG1 can efficiently seal nick DNA containing a single ribonucleotide at the 3’-end and our LIG1/DNA-RNA heteroduplex structures uncovered that LIG1 lacks a discrimination against “wrong” sugar at the final step of DNA repair (53). Yet, it remains largely unknown how the ligase active site engages with a nick DNA containing oxidative damage in the absence and presence of a ribose sugar at the 3’-end depending on the base pairing architecture, *i.e*., 8oxodG *versus* 8oxorG opposite templating A and C, remains largely unknown. These nick substrates mimic mutagenic repair intermediates that could be formed due to an incorporation of oxidized nucleotides, 8oxodGTP and 8oxorGTP opposite templating A or C, by DNA polymerases.

Base excision repair (BER), a vital guardian of the genome, is a multiprotein DNA repair pathway responsible for repairing DNA base lesions such as oxidative DNA damage (54). The BER pathway requires a coordinated function of the core repair enzymes: AP-Endonuclease 1 (APE1), DNA polymerase (pol) β, and LIG1 or LIG3α (55–57). In our previous studies, we reported that a functional interplay between polβ and the BER ligases is critical for efficient repair at the downstream steps (56–64). For example, we demonstrated that the DNA ligation step is compromised after polβ insertion of 8-oxodGTP opposite A through mutagenic base pairing, which leads to ligase failure and accumulation of abortive ligation products with a 5’-adenylate (5’-AMP) at the final steps of BER pathway (61). Moreover, we reported that polβ^+/+^ mouse embryonic fibroblast (MEF) cells display more hypersensitivity to the oxidative stress-inducing agent potassium bromate (KBrO_3_) than polβ^−/−^ cells and the *MTH1* gene deletion and the treatment of both MEF cells with an anti-cancer MTH1 inhibitor enhance this cytotoxicity to KBrO_3_ particularly in polβ^+/+^ cells (61). This supports the idea that an increased concentrations of oxidatively damaged nucleotide pools and their incorporation by polβ during BER could increase the cellular sensitivity phenotype to oxidative stress and induce ligase failure and accumulation of toxic repair intermediates with 3’-8oxoG lesion and 5’-AMP block (45–47). However, it remains undefined how a multi-protein complex involving APE1/polβ/LIG1 or LIG3α coordinate to process mutagenic nick repair intermediates containing oxidatively damaged ends at the final steps of BER pathway.

In the present study, we determined the structures of LIG1/nick DNA and DNA-RNA duplexes containing 8oxodG and 8oxorG at the 3’-strand opposite to either templating A or C. Our structures revealed that the LIG1 active site engages with the mutagenic repair intermediates containing 3’-8oxodG:A and 3’-8oxorG:A at the step 2 referring to the formation of DNA-AMP intermediate, and subsequent step 3 referring to a final phosphodiester bond formation, respectively. We also determined structures of LIG1 in complex with nick DNA harboring 3’-8oxodG:C and 3’-8oxorG:C at the second step of ligation reaction. Our structures demonstrated how the ligase active site engages with nick harboring an oxidative damage in the absence and presence of “wrong” sugar at the 3’-end through the movements at the nucleotides around nick site and by the interaction network with the critical side chains depending on dual coding potential of 8oxodGTP(*anti*):C(*anti*) and 8oxodGTP(*syn*):A(*anti*) that forms non-mutagenic Watson-Crick and mutagenic Hoogsteen base pairing, respectively, at the ligase active site.

Moreover, we showed the mutagenic nick sealing of nick DNA substrates containing 3’-8oxodG:A and 3’-8oxorG:A by LIG1 wild-type and immunodeficiency disease-associated variants *in vitro*. Our results also demonstrated an efficient ligation of 3’-8oxorG:C and a greatly diminished nick sealing of 3’-8oxodG:C by LIG1 and LIG3α. Finally, we showed a functional interplay at the downstream steps of BER pathway during an incorporation of 8oxorGTP:A by polβ coupled to subsequent ligation of resulting nick repair product by LIG1 and LIG3α in the presence of proofreading activity by APE1 for the removal of 3’-damaged base. These findings could contribute to understanding the importance of proper coordination between BER enzymes for processing of potentially toxic and mutagenic repair intermediates under the genotoxic stress conditions to maintain the repair efficiency.

## Methods

### Protein purifications

Human DNA ligase 1 (LIG1) C-terminal (△261) proteins with 6x-his tag (pET-24b) for the wild-type and mutants Glu(E)346Ala(A)/Glu(E)592Ala(A), Pro(P)529L, Arg(R)641Leu(L), and Arg(R)771Trp(W), were overexpressed and purified as reported (58–64). Briefly, the proteins were overexpressed in Rosetta (DE3) *E. coli* cells in Terrific Broth (TB) media with kanamycin (50 μgml^−1^) and chloramphenicol (34 μgml^−1^) at 37 °C. The cells were induced with 0.5 mM isopropyl β-D-thiogalactoside (IPTG) when the OD_600_ was reached to 1.0, and the overexpression was continued for overnight at 28 °C. Cells were collected in the lysis buffer containing 50 mM Tris-HCl (pH 7.0), 500 mM NaCl, 20 mM imidazole, 10% glycerol, 1 mM PMSF, and an EDTA-free protease inhibitor cocktail tablet by sonication at 4 °C. The cell lysate was pelleted at 31,000 x g for 1 hr at 4 °C. LIG1 proteins were purified by HisTrap HP column with an increasing imidazole concentration (20-300 mM) in the binding buffer containing 50 mM Tris-HCl (pH 7.0), 500 mM NaCl, 20 mM imidazole, and 10% glycerol at 4 °C. The collected fractions were subsequently loaded onto HiTrap Heparin column in the binding buffer containing 50 mM Tris-HCl (pH 7.0), 50 mM NaCl, 1.0 mM EDTA, and 10% glycerol, and then eluted with a linear gradient of NaCl up to 1 M. LIG1 proteins were further purified by Superdex 200 10/300 column in the buffer containing 20 mM Tris-HCl (pH 7.0), 200 mM NaCl, and 1 mM DTT.

Human wild-type full-length (1-922 amino acids) DNA ligase 3α (LIG3α) with 6x-his tag (pET-29a) was overexpressed in BL21(DE3) *E. coli* cells in Lysogeny Broth (LB) media at 37°C for 8 h and induced with 0.5 mM IPTG. The protein overexpression was continued overnight at 28°C. The cells were harvested, lysed at 4°C, and then clarified as described above. LIG3α protein was purified by HisTrap HP column with an increasing imidazole gradient (20-300 mM) elution at 4°C. The collected fractions were then further purified by HiTrap Heparin with a linear gradient of NaCl up to 1 and then Superdex 200 increase 10/300 column in the buffer containing 50 mM Tris-HCl (pH 7.0), 500 mM NaCl, 5% glycerol, and 1 mM DTT. DNA polymerase (pol) β (pGEX-6p-1) protein (1-335 amino acids) was overexpressed in BL21(DE3) *E. coli* cells in LB media at 37 °C for 8 h and induced with 0.5 mM IPTG. The cells were then grown overnight at 16 °C. After cell lysis at 4 °C by sonication in the lysis buffer containing 1X PBS (pH 7.3), 200 mM NaCl, 1 mM DTT, and complete protease inhibitor cocktail, the lysate was pelleted at 16,000 x rpm for 1 h and then clarified by centrifugation and filtration. The supernatant was loaded on GSTrap HP column and purified with the elution buffer containing 50 mM Tris-HCl (pH 8.0) and 10 mM reduced glutathione. To cleave GST-tag, the recombinant polβ protein was incubated with PreScission Protease for 16 h at 4 °C in the buffer containing 1X PBS (pH 7.3), 200 mM NaCl, and 1 mM DTT. After the cleavage, the polβ protein was subsequently passed through GSTrap HP column, and the protein without GST-tag was then further purified by loading onto Superdex 200 Increase 10/300 column in the buffer containing 50 mM Tris-HCl (pH 7.5), 400 mM NaCl, and 5% glycerol. AP-Endonuclease 1 (APE1) full-length (1-315 amino acids) protein (pET-24b) was overexpressed in BL21(DE3) *E. coli* cells and the cells were harvested, lysed at 4 °C, and the supernatant was loaded onto HisTrap HP column as described above. His-tag APE1 protein was purified with an increasing imidazole gradient (0-300 mM) elution and then loaded onto HiTrap Heparin column to further purify as a linear gradient of NaCl up to 1 M, and then finally loaded on Superdex 200 Increase 10/300 column in the buffer containing 50 mM Tris-HCl (pH 7.0), 500 mM NaCl, 5% glycerol, and 1 mM DTT. All proteins were concentrated and stored at −80 °C. Protein quality was evaluated by running on 10% SDS-PAGE gel and the protein concentration was determined by the A280.

### Crystallization and structure determination

LIG1 C-terminal (△261) EE/AA mutant (LIG1^EE/AA^) protein was used for crystallization as reported previously (52,53). LIG1 was mixed with the nick DNA substrates containing 8oxodG or 8oxorG at the 3’-end opposite A or C on a template position (Supplementary Table 1). LIG1 (at 27 mgml^−1^)/DNA complex solution was prepared in 20 mM Tris-HCl (pH 7.0), 200 mM NaCl, 1 mM DTT, 1 mM EDTA, and 1 mM ATP at 1.4:1 DNA:protein molar ratio and then mixed with 1 μl of reservoir solution. LIG1-nick DNA complex crystals were grown at 20 °C using the hanging drop method, harvested, and submerged in cryoprotectant solution containing reservoir solution mixed with 20% glycerol before being flash cooled in liquid nitrogen for data collection. LIG1/nick DNA complex crystals were kept at 100 °K during X-ray diffraction data collection using the beamline CHESS-7B2. X-ray diffraction data reduction and scaling were done using HKL2000 (HKL Research, Inc). All structures were solved by the molecular replacement method using PHASER with PDB entry 7SUM as a search model (65). Iterative rounds of model building were performed in COOT and the final models were refined with PHENIX (66–68). 3DNA was used for sugar pucker analysis (69). All structural images were drawn using PyMOL (The PyMOL Molecular Graphics System, V0.99, Schrödinger, LLC). Detailed crystallographic statistics are provided in Table 1.

**Table 1:**
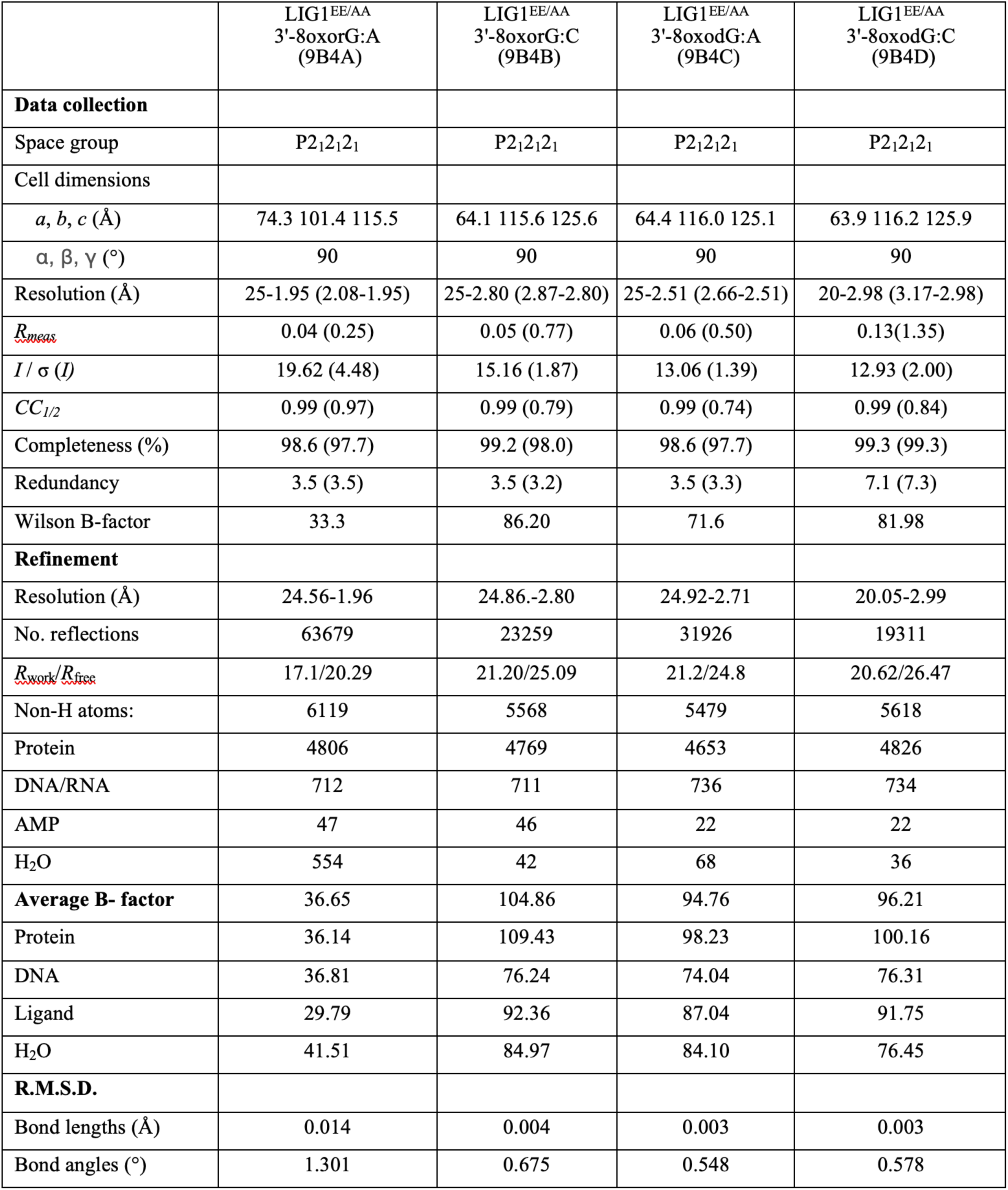
X-ray data collection and refinement statistics of LIG1 structures.

### DNA ligation assays

DNA ligation assays were performed to evaluate the ligation efficiency of LIG1 (wild-type, low-fidelity EE/AA mutant, and LIG1 deficiency disease variants P529L, R641L, R771W) as well as LIG3α (Supplementary Scheme 1) using the nick DNA substrates containing preinserted 3’-8oxodG:A, 3’-8oxodG:C, 3’-8oxorG:A, and 3’-8oxorG:C (Supplementary Table 2). In the control assays, we also tested the nick sealing in the presence of canonical DNA containing 3’-dG:C. The ligation reaction containing 50 mM Tris-HCl (pH: 7.5), 100 mM KCl, 10 mM MgCl_2_, 1 mM ATP, 1 mM DTT, 100 µgml^−1^ BSA, 1% glycerol, and nick DNA substrate (500 nM) was initiated by the addition of DNA ligase (100 nM). The reaction samples were incubated at 37 °C, stopped by quenching with an equal amount of the buffer containing 95% formamide, 20 mM EDTA, 0.02% bromophenol blue, and 0.02% xylene cyanol, and collected at the time points indicated in the figure legends. The reaction products were then separated by electrophoresis on an 18% denaturing polyacrylamide gel. The gels were scanned with a Typhoon Phosphor Imager (Amersham Typhoon RGB), and the data were analyzed using ImageQuant software.

### Nucleotide insertion assays

The insertion assays were performed to evaluate the efficiency of 8oxodGTP and 8oxorGTP insertion by polβ wild-type and Y271G mutant (Supplementary Scheme 1) using gap DNA substrates with template base A or C (Supplementary Table 3). The reaction mixture contains 50 mM Tris-HCl (pH 7.5), 100 mM KCl, 10 mM MgCl_2_, 1 mM ATP, 1 mM DTT, 100 µg ml^−1^ BSA, 1% glycerol, 8oxorGTP (100 µM), and DNA substrate (500 nM) in the final volume of 10 µl. The reaction was initiated by the addition of polβ alone (10 nM) and the reaction mixtures were incubated at 37 °C for the time points as indicated in the figure legends. The reaction products were then mixed with an equal amount of gel loading buffer, separated by electrophoresis on 18% Urea-PAGE gel, and the data were analyzed as described above.

### Coupled assays

The coupled assays were performed to evaluate the ligation efficiency by LIG1 or LIGIIIα after 8oxorGTP insertion by polβ wild-type or Y271G mutant (Supplementary Scheme 1) using gap DNA substrate with template base A (Supplementary Table 4). The reaction mixture contains 50 mM Tris-HCl (pH 7.5), 100 mM KCl, 10 mM MgCl_2_, 1 mM ATP, 1 mM DTT, 100 µg ml^−1^ BSA, 1% glycerol, 8oxorGTP (100 µM), and DNA substrate (500 nM) in the final volume of 10 µl. The reaction was initiated by the addition of the preincubated polβ/DNA ligase protein complex (100 nM) and incubated at 37 °C for the time points as indicated in the figure legends. The reaction products were then mixed with an equal amount of gel loading buffer, separated by electrophoresis on 18% Urea-PAGE gel, and the data were analyzed as described above.

### APE1 assays

The exonuclease assays were performed to evaluate the efficiency of 3’-8oxorG removal by APE1 (Supplementary Scheme 2) using nick DNA substrate containing preinserted 3’-8oxorG:A (Supplementary Table 5). The coupled assays were performed to evaluate the ligation efficiency by LIG1 or LIGIIIα after 3’-8oxorG removal by APE1 (Supplementary Scheme 2) using nick DNA substrate containing preinserted 3’-8oxorG:A (Supplementary Table 5). For both assays, the reaction mixture contains 50 mM Tris-HCl (pH 7.5), 100 mM KCl, 10 mM MgCl_2_, 1 mM ATP, 1 mM DTT, 100 µg ml^−1^ BSA, 1% glycerol, and DNA substrate (500 nM) in the final volume of 10 µl. The reaction was initiated by the addition of APE1 alone or the preincubated APE1/DNA ligase protein mixture (100 nM) and incubated at 37 °C for the time points as indicated in the figure legends. The reaction products were then mixed with an equal amount of gel loading buffer, separated by electrophoresis on 18% Urea-PAGE gel, and the data were analyzed as described above.

## Results

### Structures of LIG1/nick DNA and DNA-RNA complexes uncover the mechanism of mutagenic ligation in the presence of oxidatively damaged ends

We determined the structures of LIG1 in complex with nick DNA containing 3’-8oxodG and 3’-8oxorG opposite either Adenine (A) or Cytosine (C) on the template position (Table 1 and Figure 1) using the EE/AA mutant that harbors E346A and E592A mutations as reported previously (51–53). We solved the LIG1/DNA-RNA heteroduplex containing 3’-8oxorG:A at the step 3 of the ligation reaction referring to the formation of a phosphodiester bond between 3’-OH and 5’-PO_4_ ends of nick (Figure 1A), while LIG1/3’-8oxorG:C structure was captured at the step 2 referring to the formation of DNA-AMP intermediate when the AMP moiety is transferred to the 5’-end of the nick (Figure 1B). Similarly, we observed LIG1/nick DNA complexes containing 3’-8oxodG opposite templating A and C at the step 2 of the ligation reaction when the DNA-AMP intermediate is formed (Figure 1C-D). This step 2 structure is similar to previously reported structure of LIG1/3’-8oxodG:A (ref).

**Figure 1.**
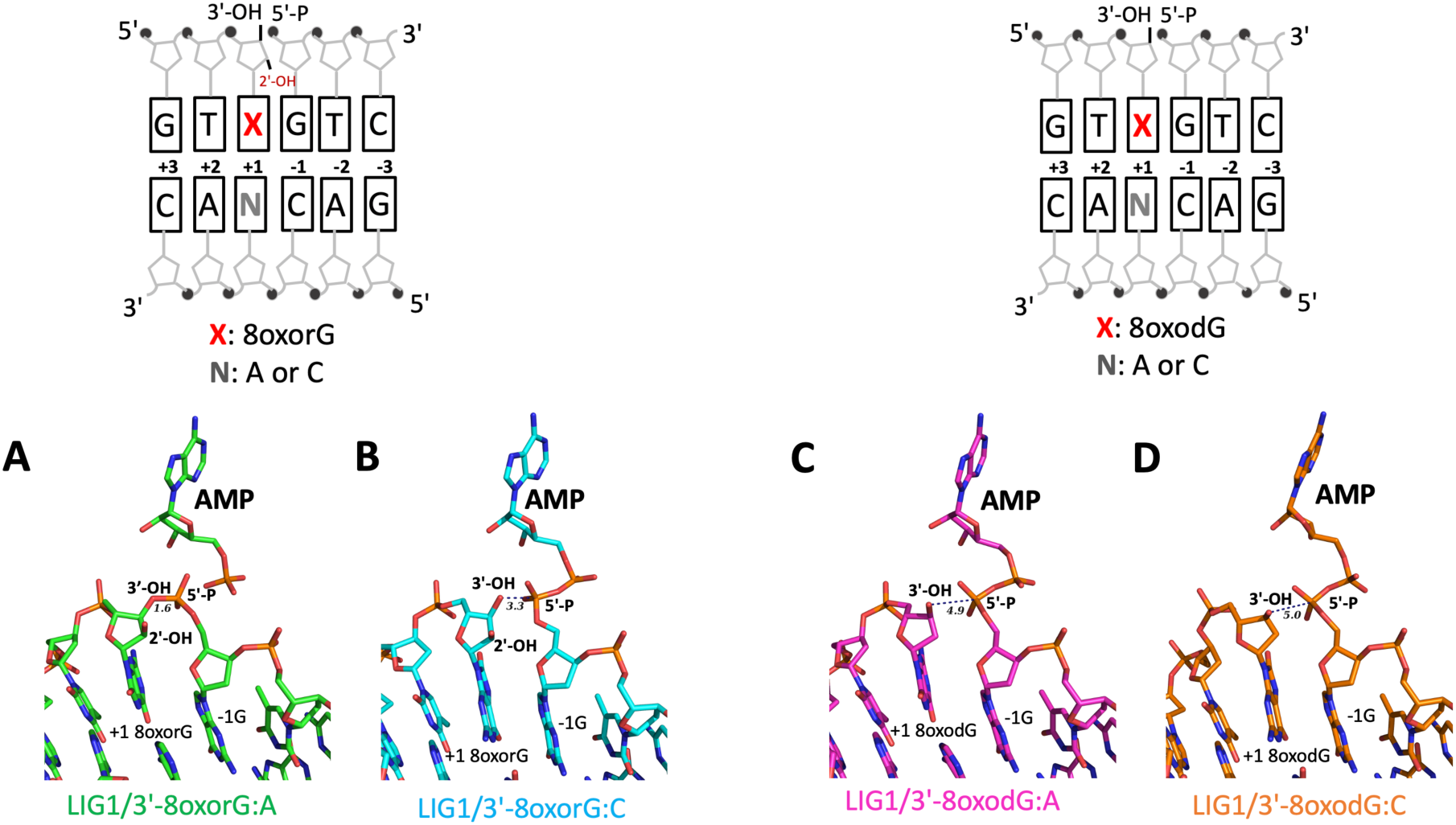
Structures of LIG1 bound to nick DNA duplexes with damaged ends. Structures of LIG1 in complex with nick DNA containing 3’-8oxorG:A (**A**), 3’-8oxorG:C (**B**), 3’-8oxodG:A (**C**) and 3’-8oxodG:C (**D**). Schematic view of DNA substrate used in the LIG1 crystallization shows the sequence of 3’- and 5’-ends at the nick site.

The omit Fo-Fc maps at 3σ demonstrate that the AMP moiety shows a continuous density for almost all the atoms in the AMP in the LIG1 step 2 structures where AMP is bound to 5’-PO_4_ of nick (Supplementary Figure 1B-D), while the omit Fo-Fc map reveals a continuous density between the 3’-OH and 5’-PO_4_ ends of the nick in the step 3 structure of LIG1/3’-8oxorG:A, demonstrating a formation of phosphodiester bond between DNA ends (Supplementary Figure 1A). The overlay of LIG1 3’-8oxodG and 3’-8oxorG structures at the step 2 *versus* 3 demonstrated differences in the position of AMP. The 3’-OH terminus attacks the 5’-PO_4_ terminus downstream of the nick DNA in the step 3 structure of LIG1/3’-8oxorG:A, while we trapped the LIG1 complex as the adenylated-DNA intermediate that is poised for nick sealing at the step 2 of the ligation reaction in the LIG1 structures of 3’-8oxodG:A, 3’-8oxodG:C and 3’-8oxorG:C (Supplementary Figure 2). Furthermore, we observed a Watson-Crick base pairing for 3’-8oxodG:C and 3’-8oxorG:C and Hoogsteen base pairing for 3’-8oxodG:A and 3’-8oxorG:A (Supplementary Figure 3). The root-mean square deviation (RMSD) of the LIG1 structures was found not higher than 0.7Å for all main chain atoms, suggesting that all LIG1 structures are in the similar conformation (Supplementary Table 6).

Furthermore, consistent with the previously solved LIG1 structures, our structures with oxidatively damaged DNA in the absence and presence of ribose demonstrate that the enzyme completely envelopes the nick DNA with the oligonucleotide-binding domain (OBD), DNA-binding domain (DBD), and adenylation domain (AdD) (Supplementary Figure 4). Sugar pucker analyses show that the ribose on 3’-8oxorG:A and 3’-8oxorG:C adopt C4’-exo and C3’-endo conformation at the 3’-end of nick, respectively, while 8oxodG adopts C1’-exo conformation (Supplementary Figure 5).

The superimposition of LIG1/nick DNA complexes containing 3’-8oxodG reveals a difference in the deoxyribose group of 8oxodG depending on the identity of templating base A or C to which 8oxoG forms Hoogsteen and Watson-Crick base pairing in -*syn* and -*anti* conformation, respectively. The overlay of both structures demonstrates a movement with the phosphodiester backbone at the +2 nucleotide relative to the 3’-OH of the nick (Figure 2A). We observed that +2T of 3’-8oxodG:C moves by 2.0Å from that of 3’-8oxodG:A (Figure 2B). Furthermore, the deoxyribose of 3’-8oxodG:C moves +2T nucleotide by 1.5Å leading to a shift by 1Å at +2A nucleotide on a template position (Figure 2C). Furthermore, the superimposition of LIG1/3’-8oxorG:A and 3’-8oxorG:C structures reveals that 8oxorG moves 0.8Å closer towards 5’-end of nick, which reduces the distance between 5’-PO_4_ and 3’-OH to 3.3Å, and both LIG1 structures share the similar interaction network with the active site residues D570 and R871 (Figure 3). These conformational changes at the nick site could be the reason for crystalizing LIG1 at the step 2 and 3 of the ligation reaction in the presence of 3’-8oxorG:C and 3’-8oxorG:A, respectively. We recently reported X-ray structures of LIG1/3’-RNA-DNA heteroduplexes containing 3’-rA:T and 3’-rG:C at the step 3 and demonstrated a lack of sugar discrimination at the ligase active site (53). The overlay previously solved LIG1/3’-rG:C structure with LIG1/3’-8oxorG:C showed that the 3’-8oxorG moves ~1Å closer towards 5’-PO_4_ of the nick leading to the 3’-OH away from the active site residue D570 but keeps the same orientation (Supplementary Figure 6).

**Figure 2.**
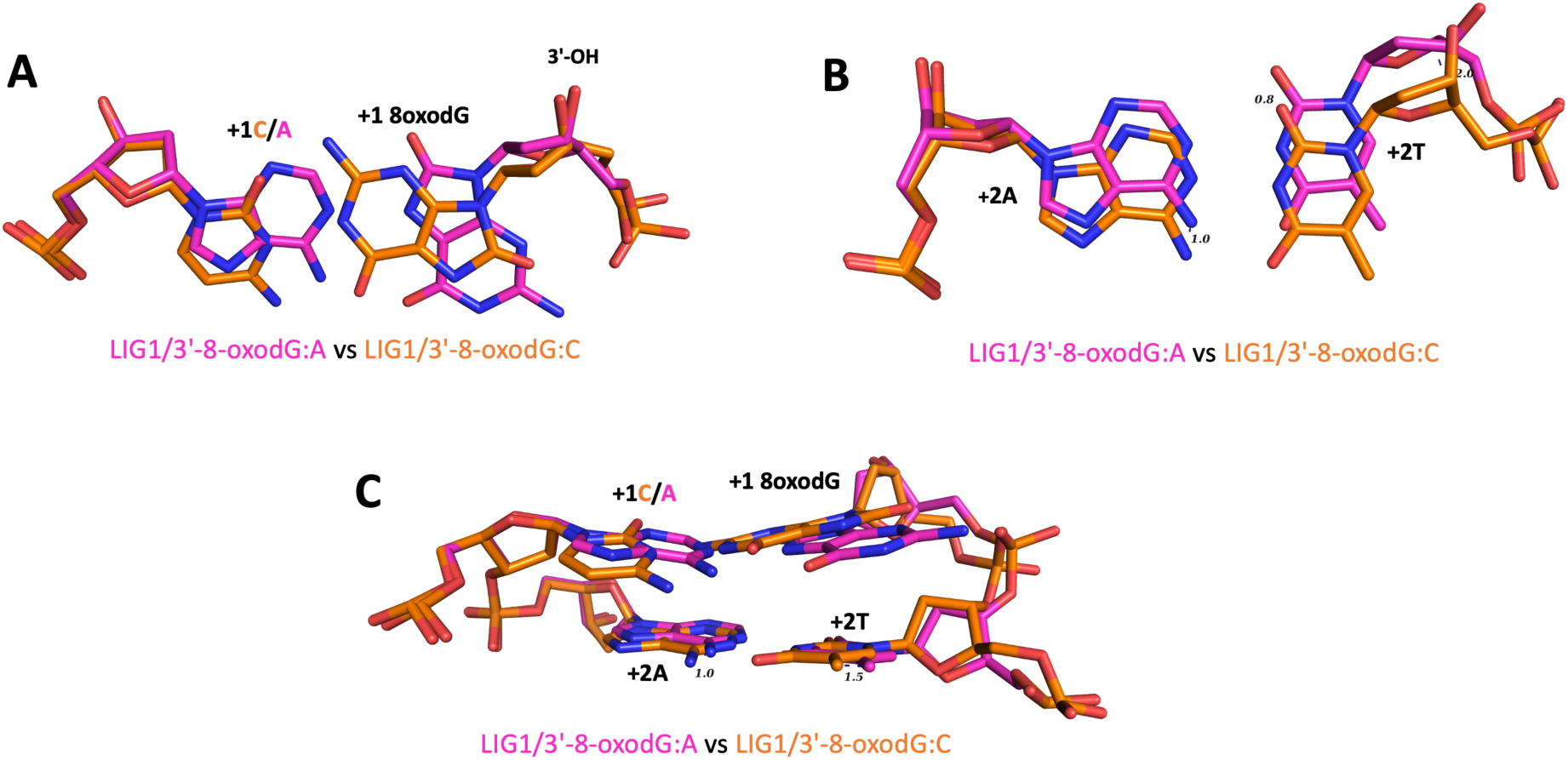
Superimposition of LIG1 structures for nick DNA complexes containing 8oxodG. The overlay of LIG1 structures demonstrates Hoogsteen base pair of 3’-8oxodG:A and Watson-Crick base pair of 3’-8oxodG:C (**A**) and movement of 1.1Å at +2T nucleotide relative to 3’-OH strand of the nick (**B**), and a conformation change at +1A/C nucleotide on templating base of 3’-8oxodG, and +2T and +2A nucleotides relative to 3’-OH strand of the nick (**C**).

**Figure 3.**
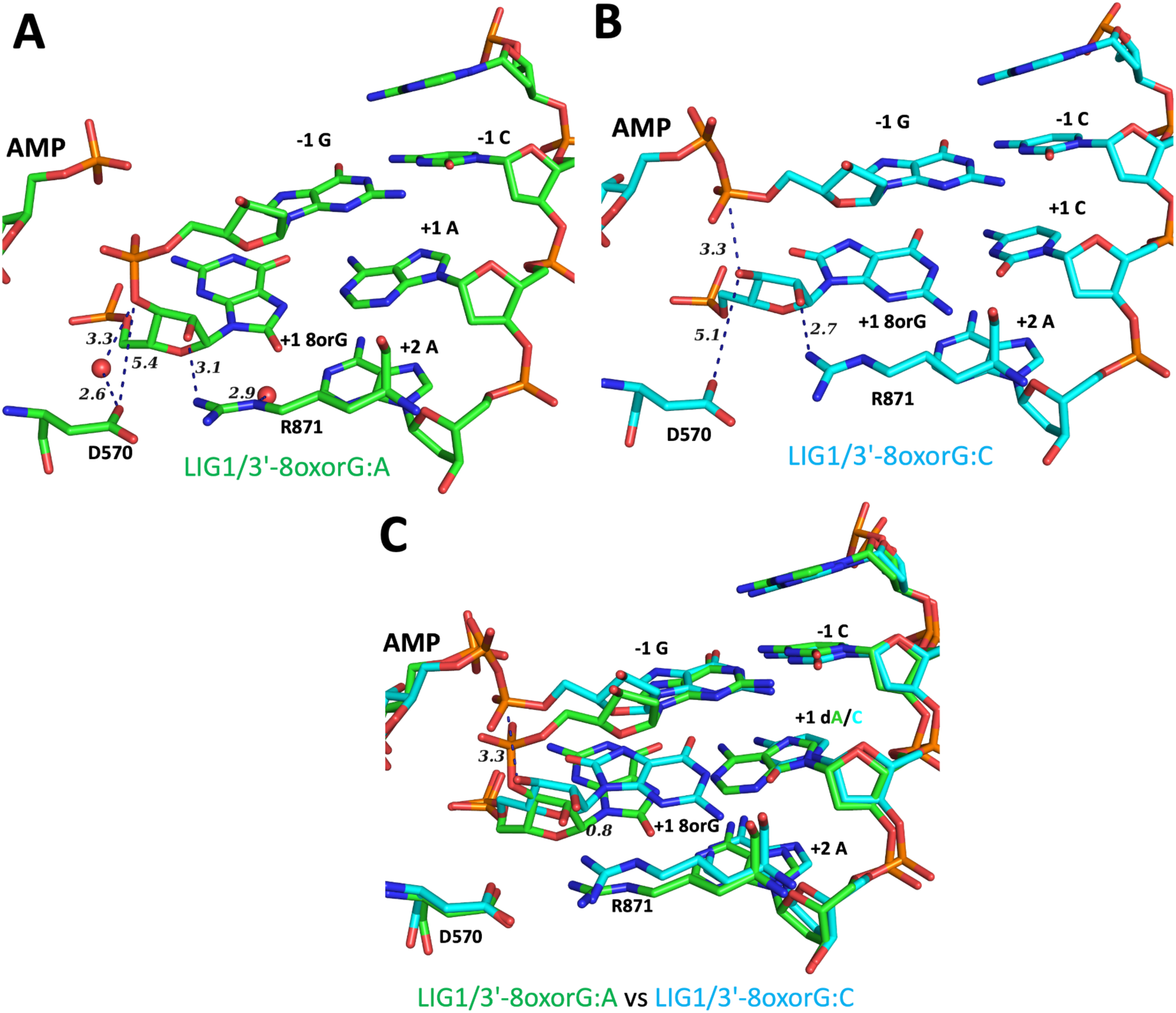
LIG1 structures of nick DNA-RNA heteroduplexes containing 8oxorG. **(A-B)** LIG1 structures of 3’-8oxorG:A (step 3) and 3’-8oxorG:C (step 2) demonstrate differences in the distances at the 3’-end of nick and the interaction network of the ligase active site residues D570 and R871. (**C**) Overlay of both LIG1 structures reveals that 8oxorG moves 0.8Å closer towards 5’-end of nick, which reduces the distance between 5’-PO_4_ and 3’-OH to 3.3Å.

### Ligation efficiency of nick DNA with oxidatively damaged ends by LIG1

We next investigated the ligation efficiency of LIG1 in the presence of the nick DNA substrates with preinserted 8oxodG and 8oxorG at the 3’-end opposite template base A or C (Supplementary Scheme 1). Our results demonstrated the mutagenic ligation of 3’-8oxodG:A and 3’-8oxorG:A by LIG1 (Figure 4A). Similarly, we observed an efficient ligation of 3’-8oxorG:C (Figure 4B, lanes 9-14), while the ability of LIG1 to seal the nick DNA substrate with 3’-8oxodG:C is greatly diminished (Figure 4B, lanes 2-7) as we reported previously (61). We did not observe a significant difference in the amount of mutagenic ligation products when 8oxodG or 8oxorG is base paired with templating A (Figure 4C). However, there was ~20-fold difference in the nick sealing products of 3’-8oxodG:C *versus* 8oxorG:C (Figure 4D). The comparisons of template base-dependent differences in the nick sealing efficiency of oxidized 3’-deoxyribo-*versus* 3’-ribo-nucleotide-containing ends demonstrated ~100-fold difference for 3’-8oxodG opposite A over C, and no difference for nick DNA containing 3’-8oxorG by LIG1 (Supplementary Figure 7).

**Figure 4.**
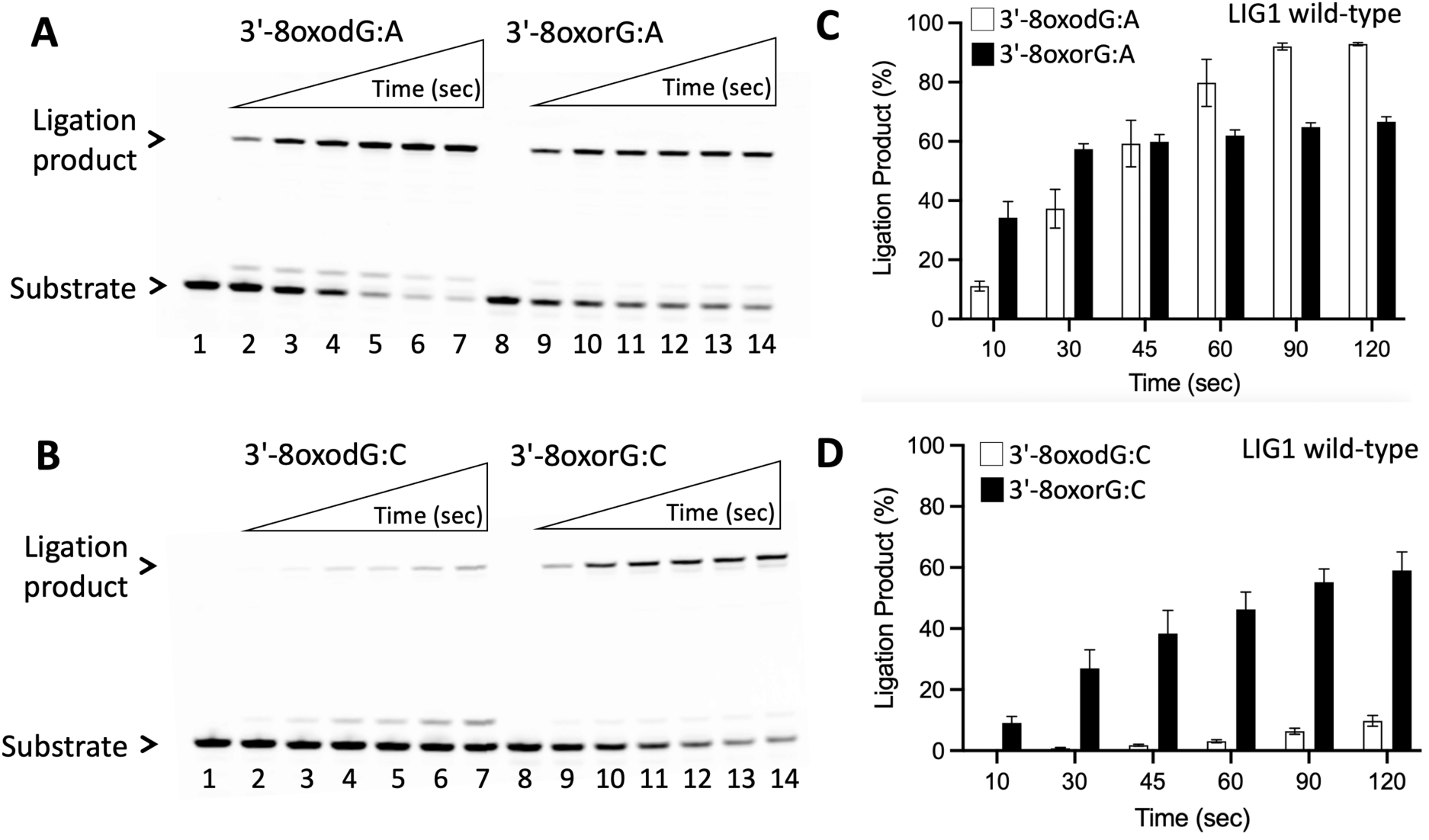
Ligation of the nick DNA substrates with damaged ends by LIG1. **(A)** Lanes 1 and 8 are the negative enzyme controls of the nick DNA substrates, and lanes 2-7 and 9-14 are the ligation products by LIG1 in the presence of 3’-8oxodG:A and 3’-8oxorG:A, respectively, and correspond to time points of 10, 30, 45, 60, 90, and 120 sec. **(B)** Lanes 1 and 8 are the negative enzyme controls of the nick DNA substrates, and lanes 2-7 and 9-14 are the ligation products by LIG1 in the presence of 3’-8oxodG:C and 3’-8oxorG:C, respectively, and correspond to time points of 10, 30, 45, 60, 90, and 120 sec. **(C-D)** Graphs show the time-dependent change in the amount of ligation products and the data represent the average from three independent experiments ± SD.

We then tested the ligation efficiency of LIG1 EE/AA low-fidelity mutant that was used in LIG1 crystallization. Our results demonstrated the mutagenic ligation of nick DNA substrates with 3’-8oxorG:A and 3’-8oxodG:A, which was similar to the efficient nick sealing of 3’-8oxorG:C (Supplementary Figure 8). We also showed a template base-dependent differences by LIG1 low-fidelity mutant EE/AA in the amount of ligation products for nick DNA substrates containing 3’-8oxodG opposite A over C and a similar efficiency for nick DNA substrates containing 3’-8oxorG:A and 3’-8oxorG:C (Supplementary Figure 9). The amount of nick sealing products by LIG1 wild-type *versus* EE/AA mutant revealed same efficiency for the oxidized ribonucleotide-containing ends, 3’-8oxorG:A and 3’-8oxorG:C (Supplementary Figure 10). As expected, in the control ligation assays, we obtained efficient nick sealing for canonical 3’-dG:C by both LIG1 wild-type and EE/AA mutant (Supplementary Figure 11).

We recently reported that LIG1 can ligate the nick DNA substrates with preinserted 3’-ribonucleotides, 3’-rA:T and 3’-rG:C, as efficient as Watson-Crick base-paired ends, 3’-dA:T and 3’-dG:C *in vitro* (53). In the present study, we next compared the ligation efficiency of nick DNA substrates with 3’-rG:C and 3’-8oxorG:C by LIG1. Our results showed more efficient ligation of nick DNA with 3’-rG:C than that of oxidatively damaged ribonucleotide-containing ends, 3’-8oxorG:C (Supplementary Figure 12A). The amount of ligation products demonstrates slightly more ligation products in the presence of 3’-rG:C than oxidatively damaged ends, especially at the earlier time points of the reaction (Supplementary Figure 12B). This could be due to conformational differences we observed in the superimposition of LIG1/3’-rG:C and 3’-8oxorG:C structures (Supplementary Figure 6).

Furthermore, we investigated the impact of mutations that have been associated with the LIG1-deficiency disease on the mutagenic nick sealing efficiency of oxidatively damaged ends (Figure 5). For this purpose, we performed ligation assays using LIG1 variants P529L, E566K, R641L, and R771W, and compared the ligation efficiency of the nick DNA substrates containing 3’-oxidized ribonucleotide. In the presence of nick DNA with 3’-8oxorG:A, when compared to the mutagenic ligation by LIG1 wild-type (Supplementary Figure 13), our results showed similar and relatively less efficient ligation with LIG1 P529L and R771W variants, respectively (Figure 5A). However, we obtained a significantly diminished nick sealing by LIG1 R641L variant with ~80-fold difference in the amount of ligation product (Figure 5B). In the presence of nick DNA with 3’-8oxorG:C, we observed efficient ligation by P529L while the ligation efficiency is completely diminished and significantly reduced by LIG1 variants R641L and R771W, respectively (Supplementary Figure 14A). The amount of ligation products demonstrated 20-to-80-dold fold difference between LIG1 variants (Supplementary Figure 14B). In the presence of canonical ends, all LIG1 variants efficiently ligate nick DNA with 3’-dG:C in the control assays (Supplementary Figure 15).

**Figure 5.**
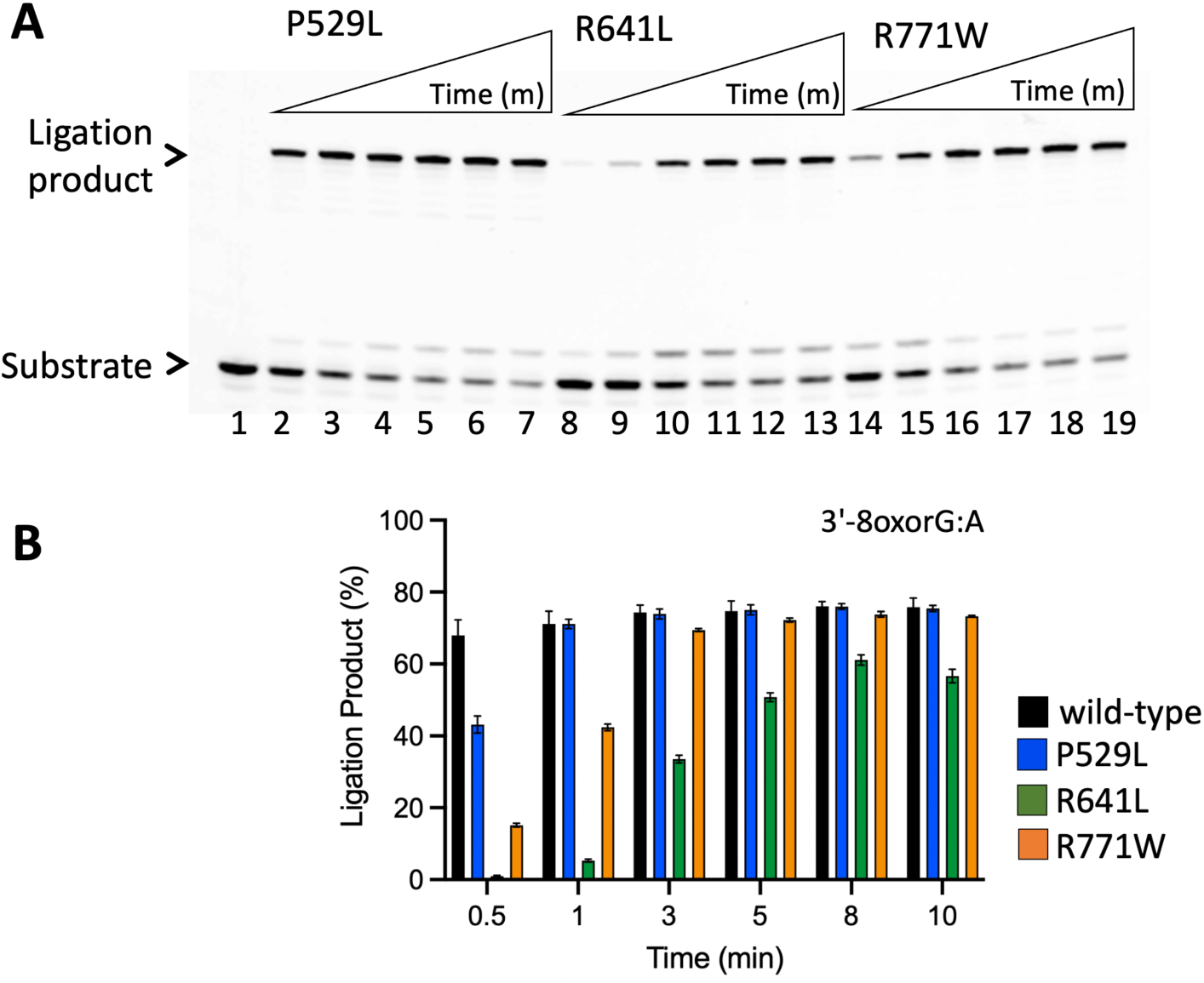
Ligation of the nick DNA substrates with damaged ribonucleotide-containing end by LIG1 deficiency disease mutants. **(A)** Line 1 is the negative enzyme control of the nick DNA substrate with 3’-8oxorG:A, and lanes 2-7, 8-13, and 14-19 are the ligation products by LIG1 variants P529L, R641L, and R771W, respectively, and correspond to time points of 0.5, 1, 3, 5, 8, and 10 min. **(B)** Graph shows the time-dependent change in the amount of ligation products and the data represent the average from three independent experiments ± SD.

### Mutagenic ligation of nick DNA with damaged ends by LIG3α

Human DNA ligases share highly conserved catalytic core and DNA-binding domain to catalyze a conserved three-step of ligation reaction using ATP and Mg^2+^ (42–44). LIG1 and LIG3α show subtle differences in the fidelity against the nick DNA substrates containing all possible 12 non-canonical mismatches at the 3’-end (59). In addition to LIG1, in the present study, we tested the ligation efficiency of LIG3α for the nick substrates containing 8oxodG or 8oxorG at the 3’-end. In the presence of the nick DNA substrate with 3’-8oxodG:A, our results demonstrated the products of mutagenic ligation and ligase failure that were appeared simultaneously (Figure 6A, lanes 2-7), while we observed products of mutagenic ligation only with 3’-8oxorG:A by LIG3α (Figure 6A, lanes 9-14). However, in the presence of nick DNA substrate with 3’-8oxodG:C, there was no ligation (Figure 6B, lanes 2-7) and an efficient nick sealing of 3’-8oxorG:C (Figure 6B, lanes 9-14) by LIG3α. The comparison of the ligation products demonstrated template base A preference of LIG3α for nick sealing of oxidatively damaged ends (Figure 6C-D). The mutagenic ligation of 3’-8oxorG:A and an efficient ligation of 3’-8oxorG:C by LIG1 *versus* LIG3α showed similar substrate specificity for oxidized ribonucleotide-containing nicks by both BER ligases (Supplementary Figures 16 and 17). However, we obtained more ligation failure products (DNA-AMP) in the presence of the nick substrates 3’-8oxodG:A and 3’-8oxodG:C by LIG3α than LIG1 (Supplementary Figure 18).

**Figure 6.**
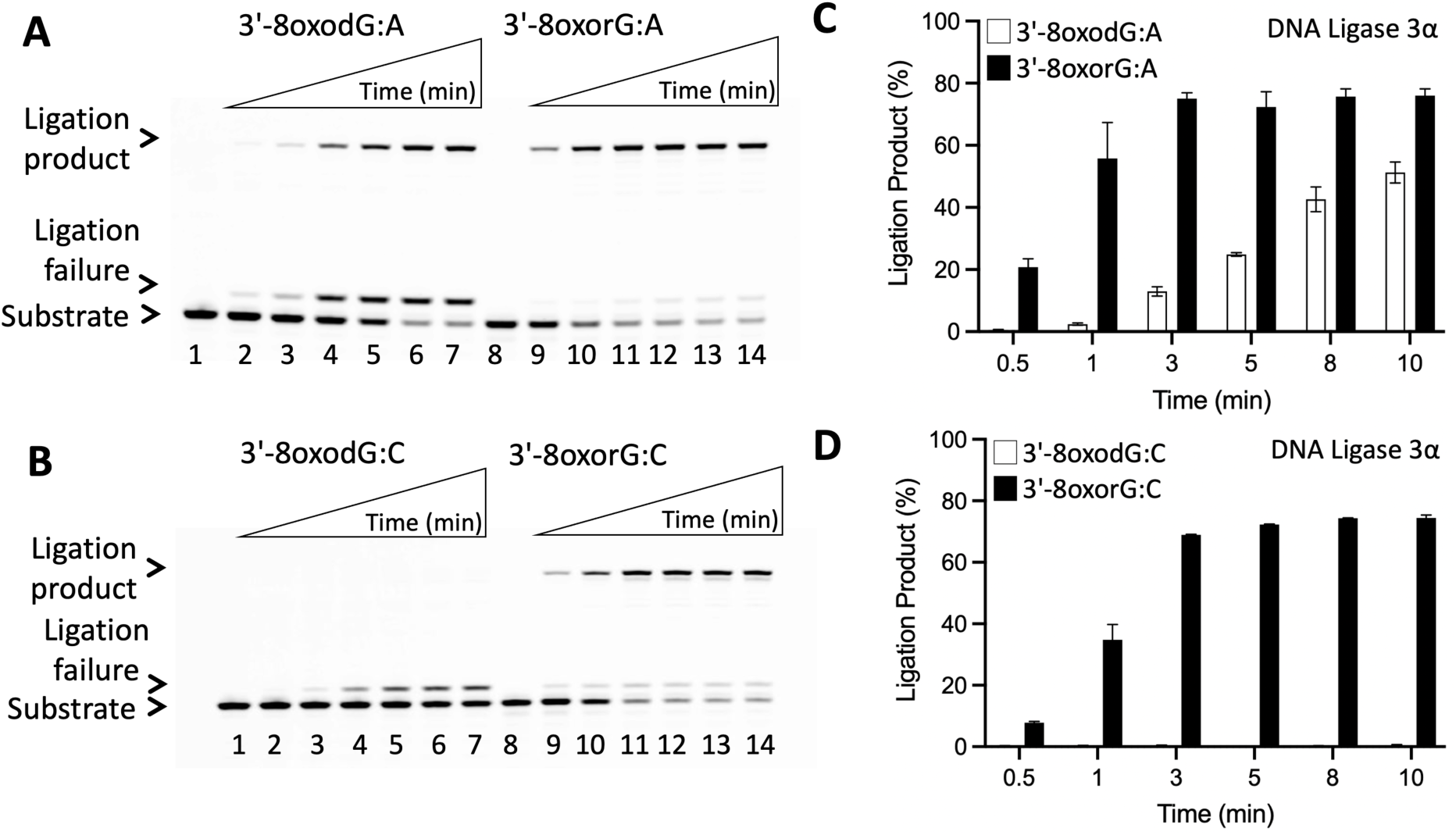
Ligation of the nick DNA substrates with damaged ends by LIG3α. **(A)** Lanes 1 and 8 are the negative enzyme controls of the nick DNA substrates, and lanes 2-7 and 9-14 are the ligation products by LIG3α in the presence of 3’-8oxodG:A and 3’-8oxorG:A, respectively, and correspond to time points of 0.5, 1, 3, 5, 8, and 10 min. **(B)** Lanes 1 and 8 are the negative enzyme controls of the nick DNA substrates, and lanes 2-7 and 9-14 are the ligation products by LIG3α in the presence of 3’-8oxodG:C and 3’-8oxorG:C, respectively, and correspond to time points of 0.5, 1, 3, 5, 8, and 10 min. **(C-D)** Graphs show the time-dependent change in the amount of ligation products and the data represent the average from three independent experiments ± SD.

### Functional coordination between APE1, polβ and BER ligases during processing of mutagenic repair intermediates at the downstream steps

We previously reported that polβ 8-oxodGTP:A incorporation compromises ligation step by LIG1 and LIG3α at the downstream steps of BER pathway leading to ligase failure products with 5’-AMP (61). As we obtained the mutagenic ligation of nick DNA with 3’-8oxorG:A by LIG1 and LIG3α (Figures 4 and 6) and solved the structure of LIG1 engaging with 3’-8oxorG:A at step 3 of ligation reaction showing a final phosphodiester bond formation (Figure 1), in the present study, we investigated the repair pathway coordination involving functions of APE1, polβ, LIG1 and LIG3α at the downstream steps in case of oxidized ribonucleotide, 8-oxorGTP, challenge (Figures 7-9).

**Figure 7.**
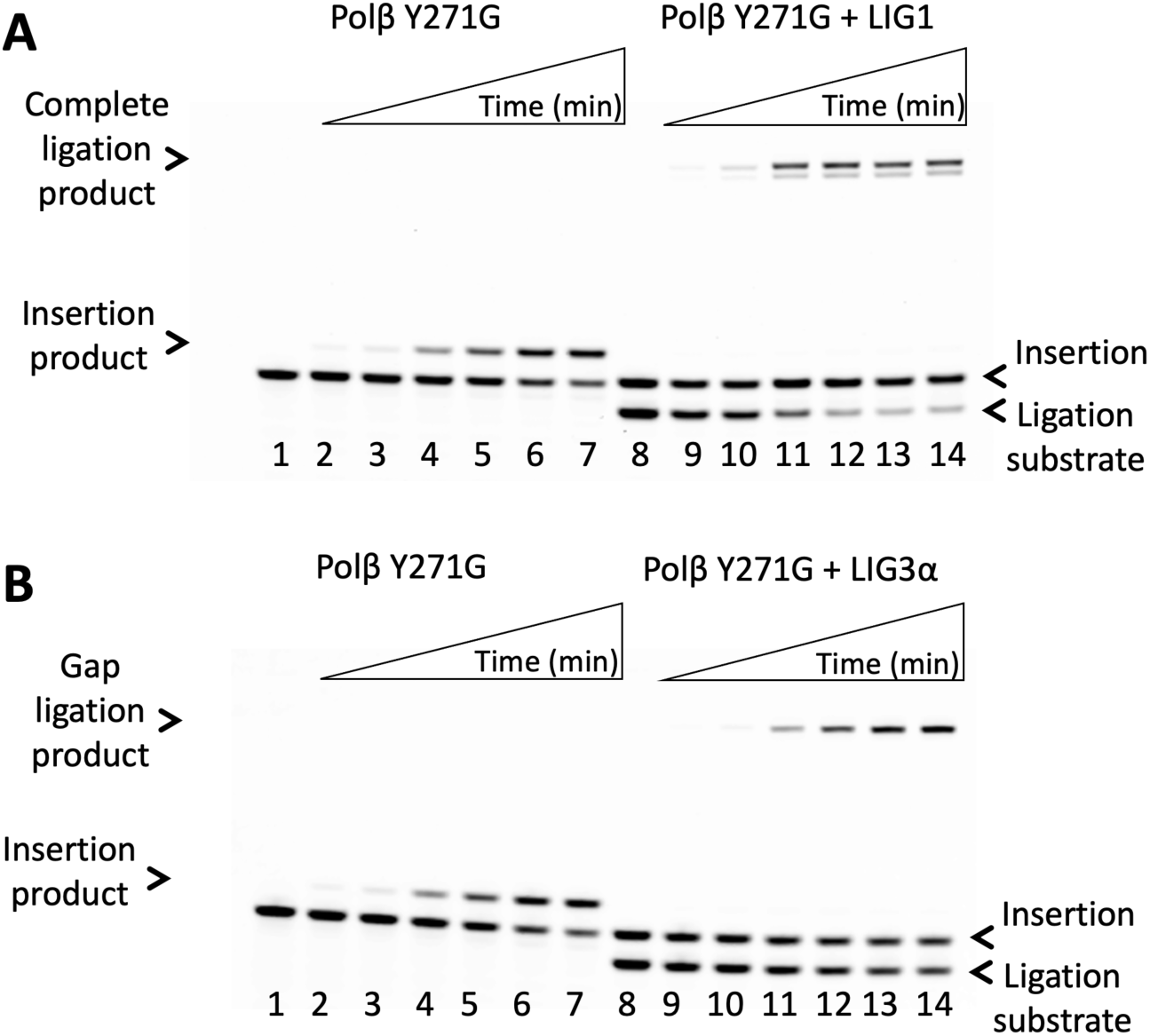
Ligation of polβ 8oxorGTP insertion products by LIG1 and LIG3α. Lanes 1 and 8 are the negative enzyme controls of gap DNA substrates for nucleotide insertion and coupled assays, respectively. Lanes 2-7 are the 8-oxorGTP:A insertion products by polβ Y271G mutant, and correspond to time points of 0.5, 1, 3, 5, 8, and 10 min. Lanes 9-14 are ligation products of polβ 8oxorGTP:A insertion by LIG1 (**A**) and LIG3α (**B**), and correspond to time points of 0.5, 1, 3, 5, 8, and 10 min.

X-ray structures of polβ reported a ribonucleotide discrimination mechanism through Tyr (Y)271 residue that acts as a steric gate to prevent the insertion and demonstrated the mutagenic insertion of 8oxorGTP at a rate similar to ribonucleotide and mismatch insertions (70,71). We first investigated the insertion efficiencies of 8oxodGTP and 8oxorGTP by polβ wild-type and Y271G mutant using one nucleotide gap DNA substrates containing A or C on templating position. Our results demonstrated similar insertion efficiency of 8oxodGTP:A by polβ and Y271G and the steric gate mutant of polβ was more efficient with 8oxorGTP:A insertion (Supplementary Figure 19). We then questioned the impact of oxidized ribonucleotide insertion by polβ on the efficiency of ligation by LIG1 and LIG3α in the same reaction mixture (Supplementary Scheme 1). Our results showed a time-dependent increase in the products of 8oxorGTP:A insertion by polβ Y271G mutant along with their conversion to nick repair intermediates and subsequent mutagenic nick sealing products by LIG1 (Figure 7A) and LIG3α (Figure 7B). The ligation of resulting nick insertion products was only observed in the coupled reaction mixture including polβ Y271G mutant that inserts 8oxorGTP:A more efficiently than wild-type. However, in the presence of polβ wild-type and DNA ligases, we mainly obtained the ligation of a gap repair intermediate itself by LIG1 and LIG3α (Supplementary Figure 20).

We recently demonstrated the role of APE1 for correcting polβ mismatch insertion products by its 3’-5’ exonuclease proofreading function while the DNA ligases attempt to seal the nick repair intermediate with non-canonical ends (60). In the present study, we also investigated the role of APE1 for proofreading nick repair intermediates containing 8oxorGTP and its coordination with BER ligases that can seal oxidatively damaged ribonucleotide-containing ends (Figure 8). Our results showed a time-dependent increase in the removal of 8oxorG from the nick DNA substrate by APE1 (Figure 8A-B, lanes 2-7). However, in the coupled reactions including APE1 and DNA ligase, the products of APE1 removal is mainly converted to gap ligation by LIG1 and LIG3α (Figure 8A-B, lanes 12-14), demonstrating that BER ligases attempt to seal resulting gap repair intermediate after the exonuclease removal of 8oxorG by APE1. We only obtained the products of mutagenic nick sealing at the earlier time points of reaction when both ligases join nick with 3’-8oxorGTP:A (Figure 8A-B, lanes 9-11)

**Figure 8.**
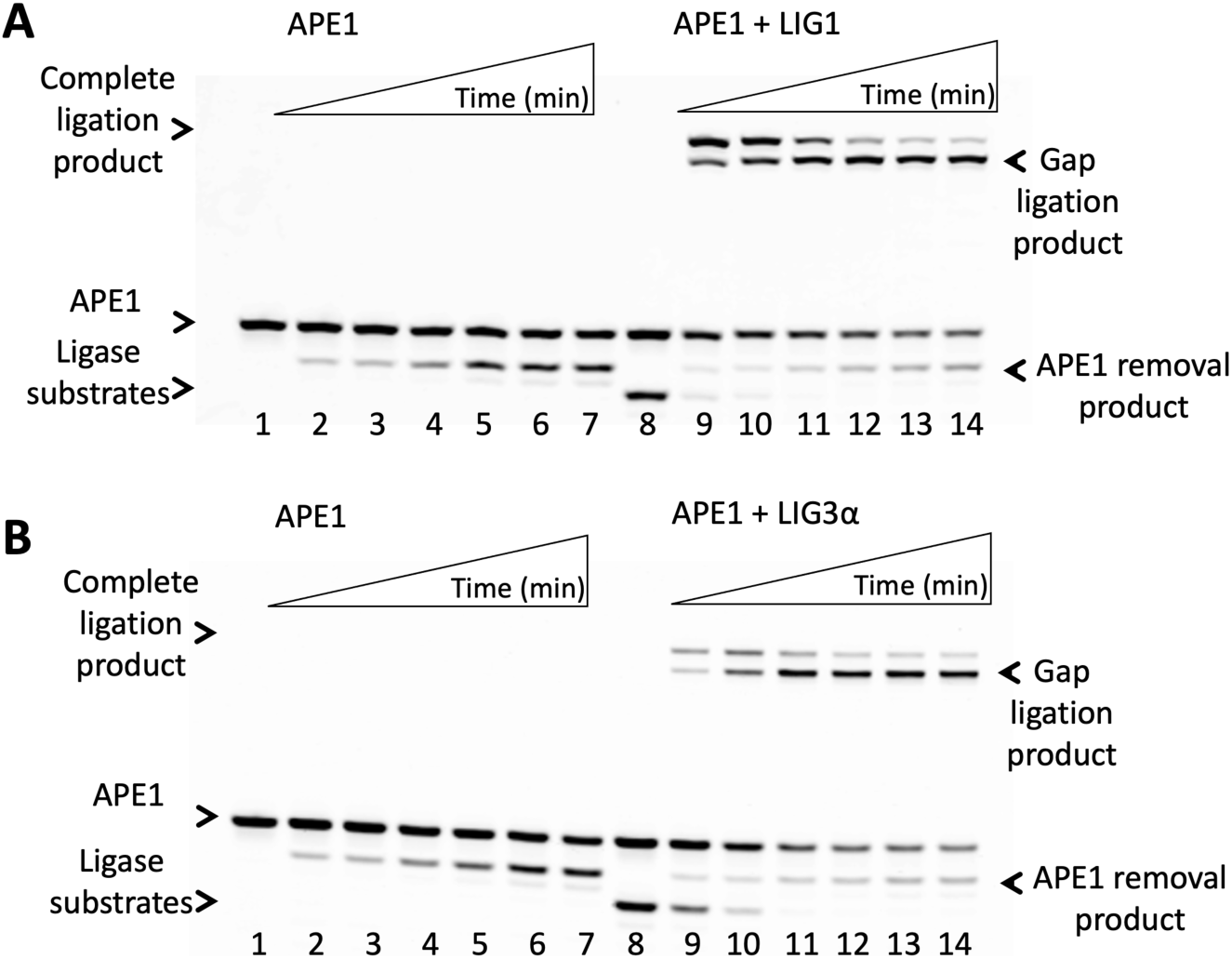
Ligation of APE1 3’-8oxorG removal products by LIG1 and LIG3α. Lanes 1 and 8 are the negative enzyme controls of nick DNA substrates for exonuclease removal and coupled assays, respectively. Lanes 2-7 are 3’-8oxorG removal products by APE1, and correspond to time points of 0.5, 1, 3, 5, 8, and 10 min. Lanes 9-14 are ligation products of APE1 3’-8oxorG removal by LIG1 (**A**) and LIG3α (**B**), and correspond to time points of 0.5, 1, 3, 5, 8, and 10 min.

## Discussion

Oxidative DNA damage plays a role in various diseases and contributes to the spontaneous mutations implicated in cancer and aging (2,3). The nucleotide pool is more vulnerable to endogenous and exogenous oxidants than chromatin-protected DNA (4). Among the nucleotides in the cellular dNTP and rNTP pools, rGTP and dGTP are the most prone to oxidation due to the high redox potential of Guanines, and their oxidation result in the formation of 8-oxo-7,8,-dihydro-2-deoxyguanosine triphosphate (8oxodGTP) and 8-oxo-guanosine-5’-triphosphate (8oxorGTP) (6). Despite expression of MTH1 that sanitizes oxidized nucleotides, very small amounts of 8-oxodGTP (<1% of the total dGTP pool as shown in mitochondria) would be enough to promote mutagenesis and drive cancer progression (7,9,20). Several DNA polymerases misincorporate 8oxodGTP in either an -*anti* or -*syn* conformation opposite to either Cytosine or Adenine, respectively, leading to A-C or G-T tranversion mutations following replication if left unrepaired (25). Furthermore, the high levels of oxidatively damaged RNA have been closely linked to aging and neurodegenerative diseases (22).

The complex network of DNA repair pathways is finalized by two consecutive steps involving a nucleotide incorporation by DNA polymerase and the subsequent rejoining of resulting nick in the DNA backbone by DNA ligase as an ultimate step (43). Upon incorporation of oxidized nucleotides by DNA polymerases with no proofreading capacity, the resulting insertion product is the nick with 3’-8oxoG lesion that could compromise ability of DNA ligase to join ends, and therefore, the ligation efficiency would be interrupted at the final step of DNA repair pathway (45–47).

Of the three human DNA ligases, LIG1 is the main replicative ligase playing a critical role for maturation of Okazaki fragments and completes most DNA repair pathways by sealing a final nick product after gap filling or a nucleotide incorporation by DNA polymerase (49). Moreover, the inherited mutations have been identified in the *LIG1* gene and reside in the C-terminal domain of the protein (P529L, E566K, R641L, and R771W), excluding the truncated polypeptide (Thr415*) that lacks the catalytic region (72). LIG1-deficiency disease patients have been mostly described with symptoms of immunodeficiency and exhibit a predisposition for cancer, stunted growth, developmental abnormalities, photosensitivity, and lymphoma (73–75). Therefore, elucidating the mechanism by which LIG1 ensures the faithful joining of DNA ends is critical for understanding how genome stability is maintained in normal and disease states.

In the present study, we structurally characterized how LIG1 engages with the potentially mutagenic and toxic repair intermediates with oxidative damage that mimic the error-prone DNA synthesis products of DNA polymerases inserting 8oxodGTP or 8oxorGTP into genome during nuclear replication and DNA repair pathways. Our structures demonstrated how the ligase active site discriminates 8oxoG at the 3’-strand, depending on the architecture of base pairing with either C or A via -*anti* or *-syn* conformation, to ensure high-fidelity of DNA ligation at the last step. We captured the ligase active site engaging with mutagenic nick containing oxidatively damaged ribonucleotide, 3’-8oxorG:A, at the step 3 of ligation reaction during which a final phosphodiester bond between 3’-OH and 5’-PO_4_ ends is formed, while our LIG1 structures of 3’-8oxorG:C was captured during step 2 when AMP is transferred to 5’-P of nick after initial ligase adenylation. Similarly, LIG1 structure in complex with nick DNA containing 3’-8oxodG:A and 3’-8oxodG:C show the step 2 of the ligation reaction referring to the formation of DNA-AMP intermediate.

We previously reported in the structures of LIG1/non-canonical nick complexes that the active site exhibits distinct conformational changes depending on the primer:terminus base pairing (A:C or G:T mismatches), and lacks a sugar discrimination against a single ribonucleotide (rA:T and rG:C) at the 3’-end (52,53) (Supplementary Table 7). It’s important to note that the previously solved structure of LIG1 bound to oxidatively damaged DNA, 8oxodG:A, by other group revealed that the ligase employs metal-reinforced DNA binding to validate base pairing via a conserved Mg^2+^-binding site at 3’-strand of nick and ligase fidelity is governed by a high-fidelity interface between LIG1, Mg^2+^, and the DNA substrate (51). Overlay of both LIG1/3’-8oxodG:A structures showed no significant difference in the active site interactions at nick site (Supplementary Figure 21).

In the ligation assays *in vitro*, we showed the mutagenic ligation of nick DNA substrates with 3’-8oxodG:A and 3’-8oxorG:A, and an efficient nick sealing of 3’-8oxorG:C, while there was a greatly compromised end joining of 3’-8oxodG:C by LIG1 (Figure 4 and Supplementary Figure 7). The overlay of LIG1/3’-8oxodG:A *versus* LIG1/3’-8oxodG:C structures demonstrated a conformational change at the deoxyribose group of 8oxodG and a significant movement at +2T and templating +2A nucleotides relative to 3’-OH strand of nick. This could be the reason for diminished nick sealing ability of LIG1 in the presence of nick DNA substrate with 3’-8oxodG:C in comparison with efficient ligation of all other nick substrates we observed (Figure 4). The superimposition of these structures with our previously solved structure of LIG1/3’-dG:C further provides an insight into this movement at the nick (Supplementary Figure 22). The overlay of LIG1/3’-dG:C and 3’-8oxoG:A suggests that a small movement at the +2 nucleotide (0.4Å and 0.7Å at +2A and +2T, respectively) has no impact on the overall movement on 3’-8oxoG:A. However, in the overlay of LIG1/3’-dG:C and 3’-8oxoG:C, the movement at the +2 nucleotides relative to 3’-OH strand of nick (1.4Å and 2.2Å at +2A and +2T, respectively) causes a significant displacement in the opposite direction to the active site. Furthermore, 3’-8oxodG:A in *syn*-conformation provides more flexibility at 3’-8oxodG towards 5’-PO_4_ of nick. Yet, 8oxodG:C in -*anti*-conformation and stable base paring at the nick restrains the movement of 3’-8oxodG towards 5’-PO_4_ of nick. Although we observed an efficient ligation of nick DNA substrates in the presence of oxidized ribonucleotide either templating A or C with similar amount of ligation products (Supplementary Figure 7), the overlay of LIG1/3’-8oxorG:A *versus* 3’-8oxorG:C (Figure 3) structures shows the significant displacement (0.8Å) at the 3’-8oxorG towards 5’-end of the nick in the LIG1/3’-8oxorG:C (step 2 structure) with reference to LIG1/3’-8oxorG:A (step 3 structure) and the differences in the interaction network between 2’-OH of 3’-8oxorG and the side chain R871. This causes conformational changes at the nick and could be the reason for capturing the crystal structures of LIG1/3’-8oxorG:A and 3’-8oxorG:C at different steps of the ligation reaction. We obtained similar ligation profile of nick substrates containing 8oxorG templating either A or C by LIG3α although there was more ligase failure products of DNA-AMP and less efficient ligation of 8oxodG templating either A or C than those of LIG1 (Figures 4 and 6). Moreover, we showed mutagenic nick sealing of 3’-8oxorG:A by LIG1-deficiency disease mutants with relatively less efficiency than that of LIG1 (Figure 5). Further structure/function studies with LIG1 variants and LIG3α with nick complexes containing oxidatively damaged DNA with and without ribose sugar are required to better understand the mechanism of faithful ligation by human DNA ligases at the final steps of DNA repair pathways.

Finally, in our study, we investigated a functional interplay between downstream enzymes of BER pathway, APE1, polβ, LIG1, and LIG3α, to understand the processing of mutagenic repair intermediates in the presence of oxidized ribonucleotides. The steps of the BER pathway involve the coordination between repair enzymes to receive the DNA substrate and efficiently pass the resulting repair product along to the next enzyme in a sequential manner, which could prevent the accumulation of potentially toxic and mutagenic DNA strand break intermediates that could trigger cell cycle arrest, necrotic cell death, or apoptosis as well as harmful nuclease activities or recombination events (56,57). During the downstream steps of the BER pathway, polβ incorporates a nucleotide into gap repair intermediate, which generates a nick product for the next enzyme in the BER pathway where LIG1 or LIG3α ultimately joins adjacent 3’-OH and 5’-PO_4_ ends to complete the repair process at the final step of BER pathway (47). Using time-lapse crystallography, the structural characteristics of polβ with an incoming 8-oxodGTP revealed how the polymerase active site discriminates between damaged and non-damaged nucleotides and the insertion can be accommodated in either the *syn*- or *anti*-conformation (27). Furthermore, the structural snapshots of reaction intermediates demonstrated that, with either templating base A or C, hydrogen-bonding interactions between the bases are lost and the enzyme reopens after catalysis, leading to a problematic nick repair intermediate (27). Similarly, X-ray structures of polβ reported that the polymerase can insert 8oxorGTP opposite to either template base or discriminates against ribonucleotides through a steric gate residue, Tyr(Y)271, to prevent the insertion of ribonucleotides (71). In our previous study, we further investigated the impact of this resulting mutagenic nick insertion product on the next ligation step of the BER pathway and reported that both LIG1 and LIG3α fail on that repair intermediate leading to the accumulation of ligation failure products with 3’-8oxoG lesion and 5’-AMP block (61). As a fidelity checkpoint of BER, it has been demonstrated that APE1 proofreads polβ-mediated errors by removing the mis-inserted bases by its 3’-5’ exonuclease activity (76). Similarly, the studies using purified proteins and human cell extracts *in vitro* reconstituted system demonstrated that the repair of 3’-8oxoG is resistant to excision by DNA glycosylases involved in the repair of oxidative lesions and APE1 can remove 3’-8oxoG lesions when incorporated by DNA polymerases during repair (76). In our previous study, we reported that APE1 can coordinate with both polβ and BER ligases during an exonuclease removal of mismatches from the nick repair intermediate with non-canonical end while LIG1 and LIG3α can attempt to seal it at the final steps of the repair pathway (60). Although it has been well characterized biochemically and at atomic level that how polβ and APE1 can process oxidative DNA damage at earlier steps, the functional coordination of both BER proteins with the downstream ligases, LIG1 and LIG3α, during processing of mutagenic repair intermediates with damaged ends at the final steps remains unknown. Our results demonstrated that both ligases can seal nick repair product of polβ 8oxorGTP:A incorporation while APE1 can serve as a fidelity check-point by removing 8oxorG. We also showed that APE1 can coordinate with LIG1 or LIG3α on the nick repair intermediate containing 8oxorG:A during the removal of 3’-oxidized base removal *versus* the ligation of resulting gap repair intermediate resulting in deleterious DNA intermediates. These findings could contribute to understanding how a multi-protein BER complex coordinate at the downstream steps during the processing of mutagenic repair intermediates.

Overall findings could provide insight into the ligase/nick DNA interactions in the presence of mutagenic nick with an oxidative damage after an incorporation of oxidized nucleotides by DNA polymerases during DNA repair and replication, which is critical given the role of oxidized nucleotides in mutagenesis, cancer therapeutics, and bacterial antibiotics. Our structures could contribute to the understanding of how LIG1 ensures fidelity depending on the dual coding potential of 8oxodGTP(*anti*):C and 8oxodGTP(*syn*):A at the ligase active site. Insights from our structures could uncover novel molecular information on the ramifications of mutagenic nick sealing by LIG1, which is critical to understand disease mechanisms, cancer development, and treatment effects. Understanding of the molecular determinants that dictate ligation fidelity at the final step of almost all DNA repair pathways could expose breakthrough platforms to manipulate the oxidative DNA damage response for future therapeutic targeting in cancer cells with higher level of genotoxic stress.

## Data and material availability

Atomic coordinates and structure factors for the reported crystal structures of LIG1 have been deposited in the RCSB Protein Data Bank under accession numbers 3’-8oxodG:A (9B4C), 3’-8oxodG:C (9B4D), 3’-8oxorG:A (9B4A), and 3’-8oxorG:C (9B4B). All data are contained within the manuscript. Further information and requests of materials used in this research are available from the authors upon reasonable request and should be directed to Dr. Melike Çağlayan.

## Acknowledgement

This work is based upon research conducted at the Center for High Energy X-ray Sciences (CHEXS), which is supported by the National Science Foundation under award DMR-1829070, and the Macromolecular Diffraction at CHESS (MacCHESS) facility, which is supported by award 1-P30-GM124166-01A1 from the National Institute of General Medical Sciences, National Institutes of Health, and by New York State’s Empire State Development Corporation (NYSTAR).

