## Supplementary Data for "Structural and biochemical characterization of LIG1 during mutagenic nick sealing of oxidatively damaged ends at the final step of DNA repair"

Supplementary Figures 1-22

Supplementary Tables 1-7

Supplementary Schemes 1-2

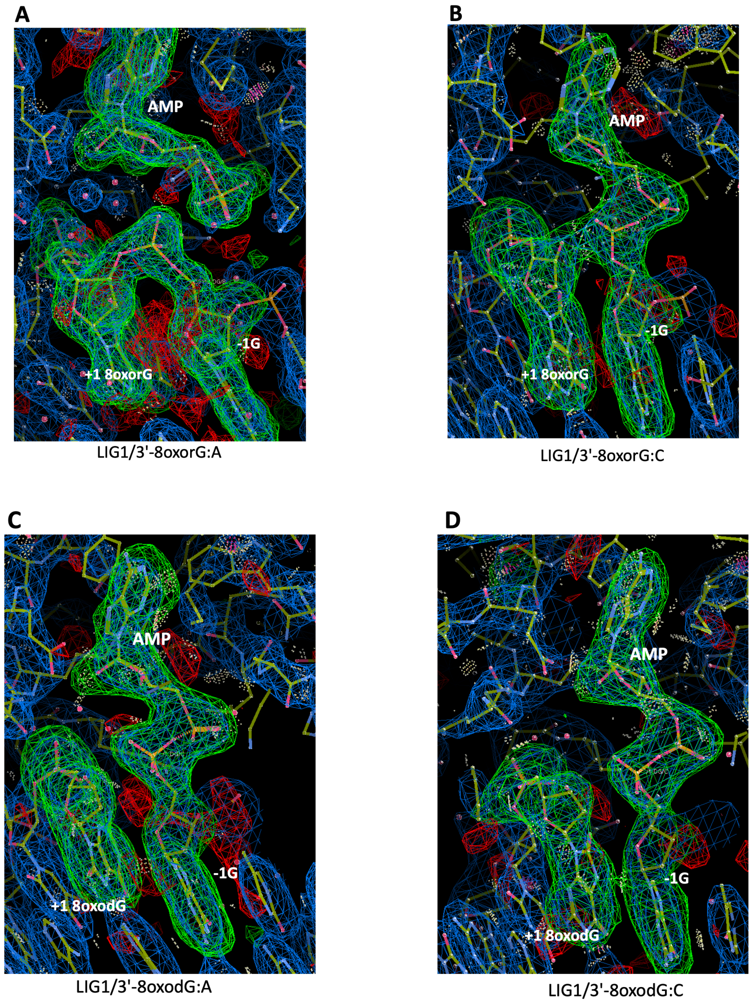


**Supplementary Figure 1. Structures of LIG1/nick DNA complexes show differences at the position of AMP. (A-B)** Structures of LIG1 in complex with oxidatively damaged DNA/RNA heteroduplexes were shown for the step 3 structure of 3'-8oxorG:A and the step 2 structure of 3'-8oxorG:C. **(C-D)** Structures of LIG1 in complex with oxidatively damaged DNA complexes were shown for the step 2 structures of 3'-8oxodG:A and 3'-8oxodG:C. All step 2 structures contain AMP bound to 5'-$PO_4$ end of nick. Simulated annealing omit maps (Fo-Fc) of the AMP are contoured at 3σ.

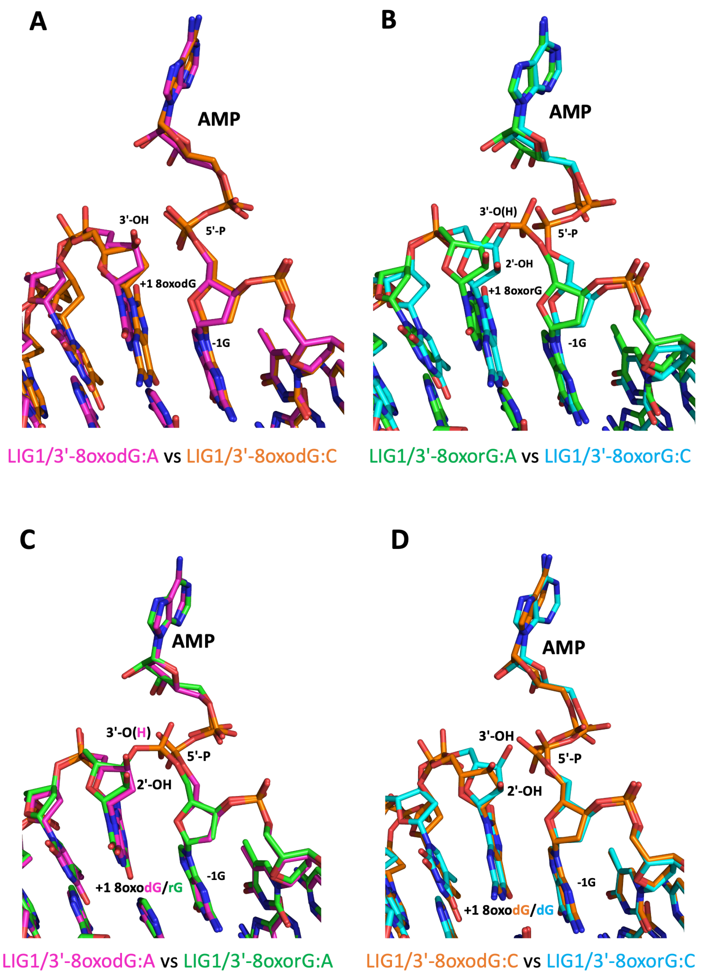


**Supplementary Figure 2. Overlay of LIG1 DNA and DNA/RNA heteroduplexes containing oxidatively damaged ends. (A)** Overlay of 3'-8oxodG:A and 3'-8oxodG:C shows that 3'-OH of the nick shares the same positions. **(B)** Overlay of 3'-8oxorG:A and 3'-8oxorG:C show that the ribose sugar at the 3'-end of the nick shares the similar conformation, yet it moves closer toward the 5'-end in the step 3 structure of 3'-8oxorG:A. (**C**-**D**) Overlays of 3'-8oxodG:C and 3'-8oxorG:C and 3'-8oxodG:A and 3'-8oxorG:A shows that 3'-8oxoG adopts similar orientation and conformational changes at the 3'-OH of nick.

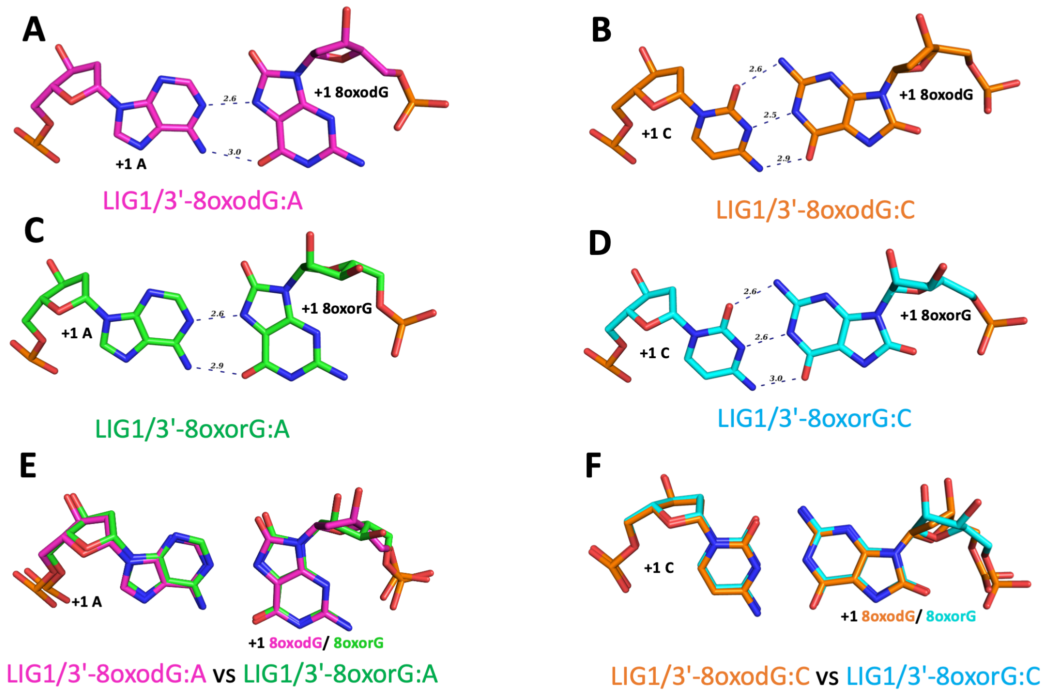


**Supplementary Figure 3. Base-pairing architecture of LIG1 structures. (A-D)** Hoogsteen and Watson-crick base pairings at the 3'-terminus of the nick are shown for the structures of LIG1/nick complexes 3'-8oxodG:A and 3'-8oxorG:A as well as 3'-8oxodG:C and 3'-8oxorG:C, respectively. **(E-F)** The overlay of LIG1/nick DNA complex structures for Hoogsteen base pairs (E, 3'-8oxodG:A and 3'-8oxorG:A) and Watson-Crick base pairs (F, 3'-8oxodG:C and 3'-8oxorG:C) showsn o difference in base-pairing architecture.

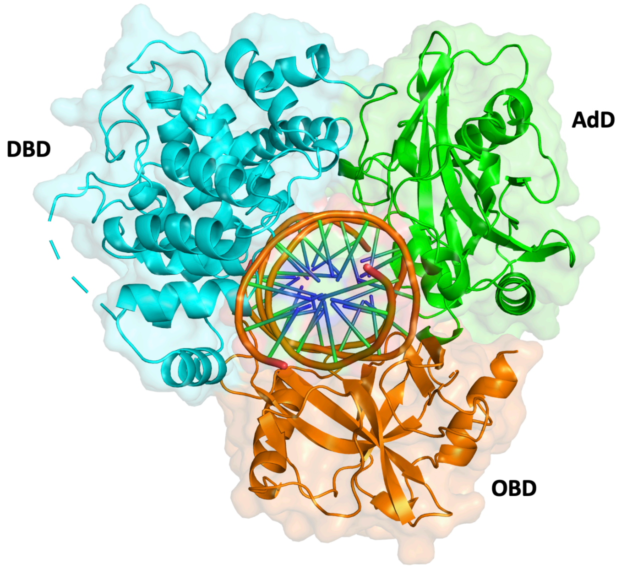


**Supplementary Figure 4. X-ray structures of LIG1/nick DNA complexes.** The superimposition of all four LIG1 structures in the present study demonstrates a global conformation that the catalytic core adopts with oxidatively damaged ends. LIG1 catalytic core consists of Adenylation (AdD, green), DNA-binding (DBD, cyan), and Oligonucleotide-binding (OBD, blue) domains that encircle the nick DNA substrate (orange).

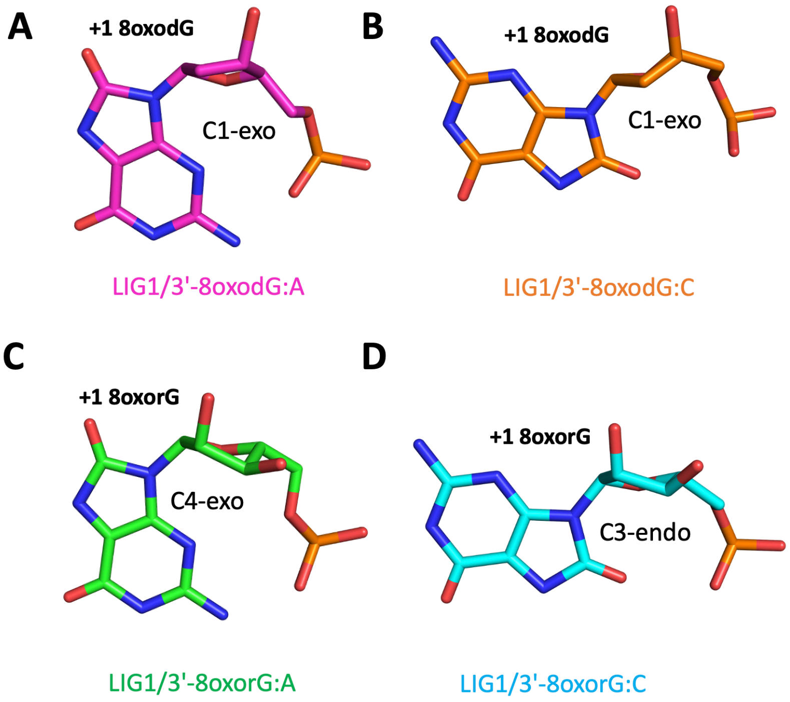


**Supplementary Figure 5. Sugar pucker analyses of LIG1 structures.** 3'-end sugar pucker analyses of LIG1 structures are presented for nick DNA complexes containing 3'-8oxodG:A (**A**), 3'-8oxodG:C (**B**), 3'-8oxorG:A (**C**), and 3'-8oxorG:C (**D**).

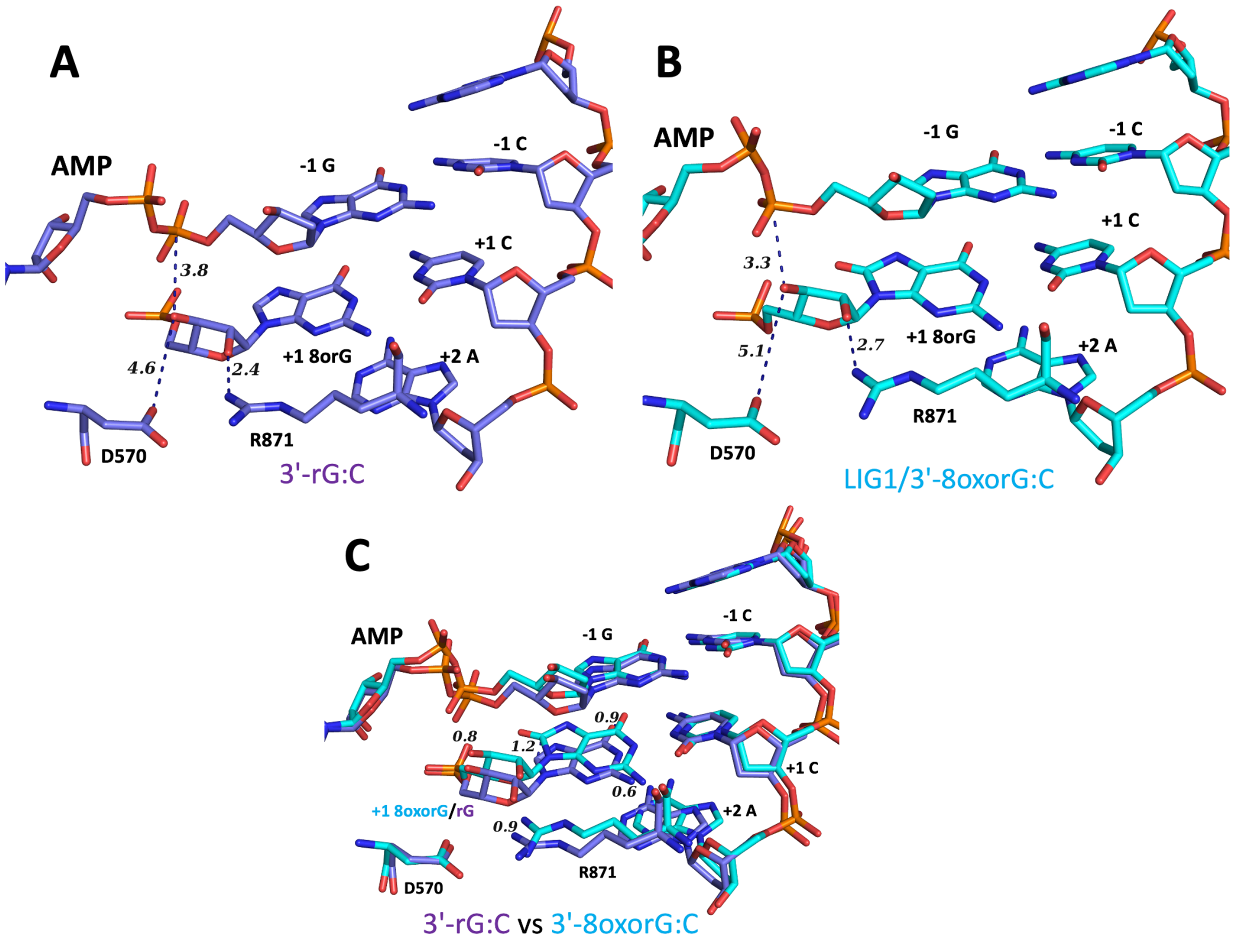


**Supplementary Figure 6. Overlay of LIG1/RNA-DNA heteroduplexes with and without oxidatively damaged ribonucleotide.** (**A-B**) LIG1 structures of 3'-rG:C and 3'-8oxorG:C demonstrate differences at the 3'- and 5'-ends of nick as well as the interaction network with the ligase active site residues D570 and R871. (**C**) Overlay of both LIG1 structures shows that the 3'-8oxorG:C adopts similar conformation with 3'-rG:C at the nick site. The 3'-end of the +1C nucleotide in the structure of 3'-8oxorG:C moves closer to the 5'-end resulting in a movement at 5'-P of -1G nucleotide relative to nick site. We previously solved LIG1/3'-rG:C structure (PDB:8VZL,53)

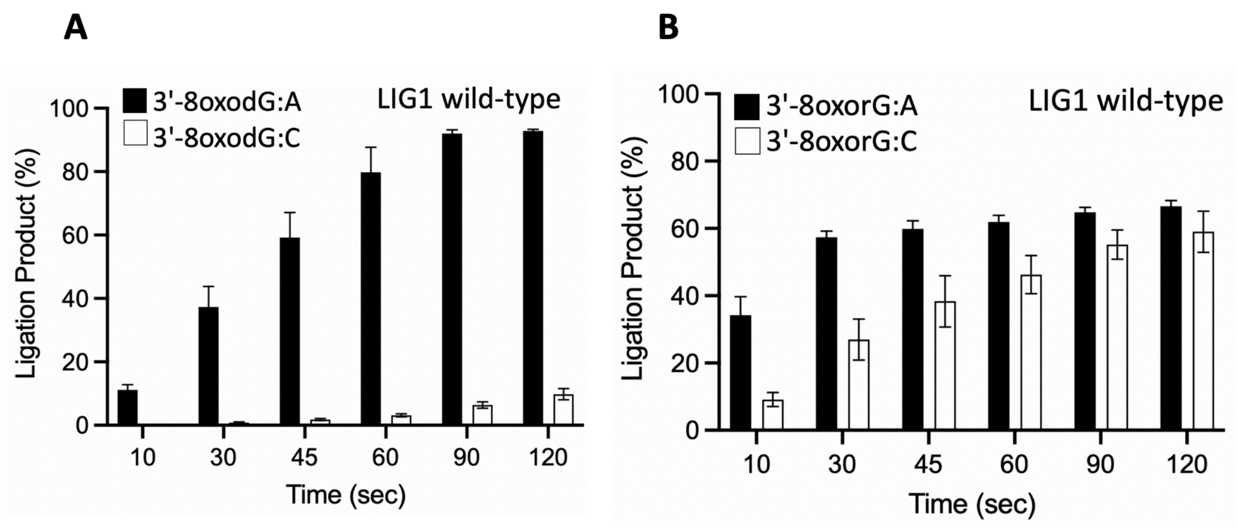


**Supplementary Figure 7. Ligation of the nick DNA substrates with damaged ends by LIG1.** Graphs show the time-dependent change in the amount of ligation products for nick DNA substrates containing 3'-8oxodG:A *versus* 3'-8oxodG:C (**A**) and 3'-8oxorG:A *versus* 3'-8oxorG:C (**B**) by LIG1 wild-type. The data represent the average from three independent experiments ± SD.

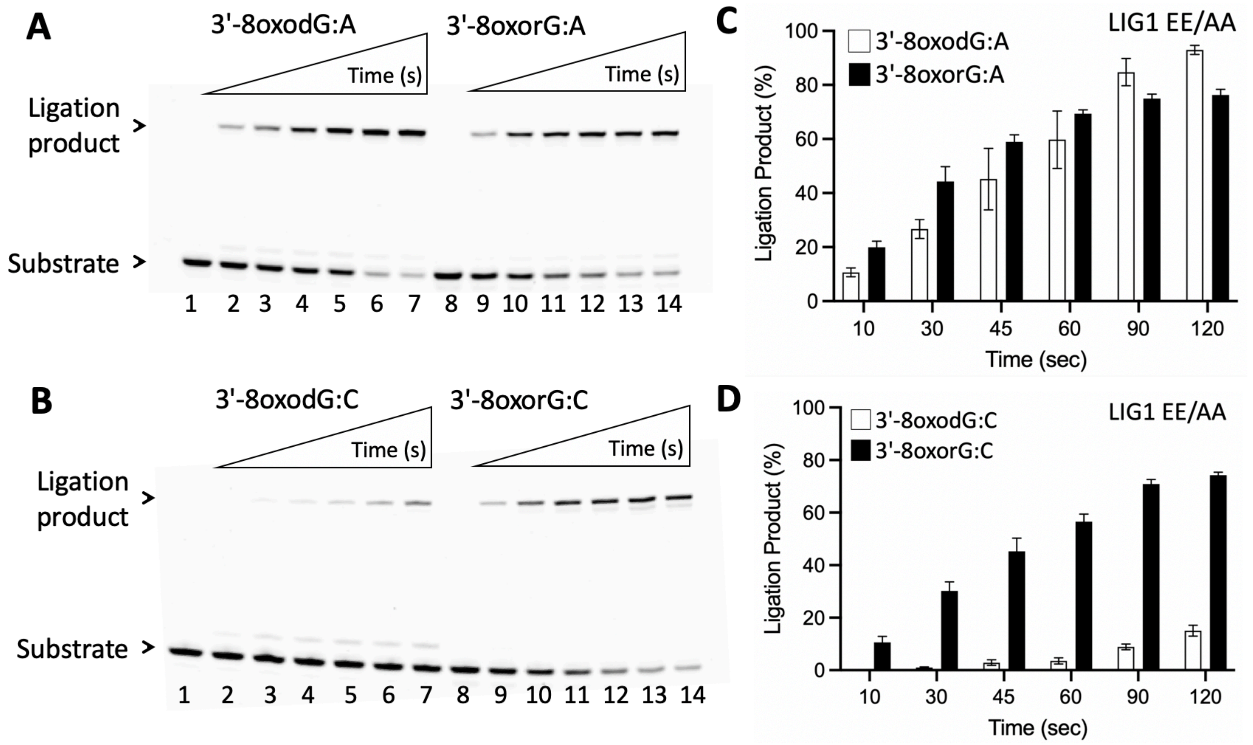


**Supplementary Figure 8. Ligation of the nick DNA substrates with damaged ends by LIG1 low-fidelity mutant EE/AA. (A)** Lanes 1 and 8 are the negative enzyme controls of the nick DNA substrates, and lanes 2-7 and 9-14 are the ligation products by LIG1 EE/AA mutant in the presence of 3'-8oxodG:A and 3'-8oxorG:A, respectively, and correspond to time points of 10, 30, 45, 60, 90, and 120 sec. **(B)** Lanes 1 and 8 are the negative enzyme controls of the nick DNA substrates, and lanes 2-7 and 9-14 are the ligation products by LIG1 EE/AA mutant in the presence of 3'-8oxodG:C and 3'-8oxorG:C, respectively, and correspond to time points of 10, 30, 45, 60, 90, and 120 sec. **(C-D)** Graphs show the time-dependent change in the amount of ligation products and the data represent the average from three independent experiments ± SD.

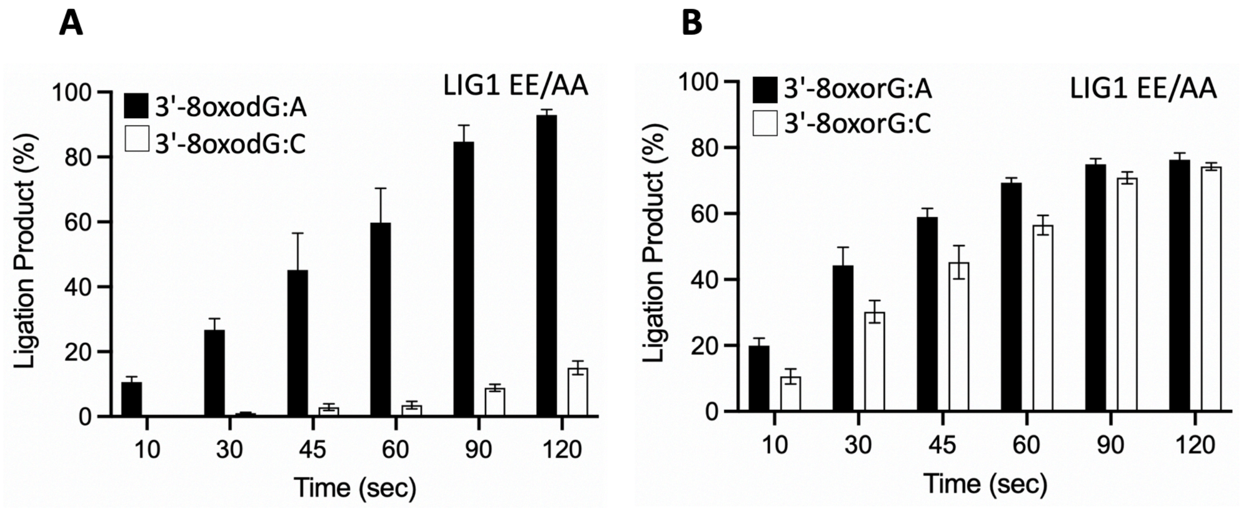


**Supplementary Figure 9. Ligation of the nick DNA substrates with damaged ends by LIG1 EE/AA.** Graphs show the time-dependent change in the amount of ligation products for nick DNA substrates containing 3'-8oxodG:A *versus* 3'-8oxodG:C (**A**) and 3'-8oxorG:A *versus* 3'-8oxorG:C (**B**) by LIG1 low-fidelity mutant EE/AA. The data represent the average from three independent experiments ± SD.

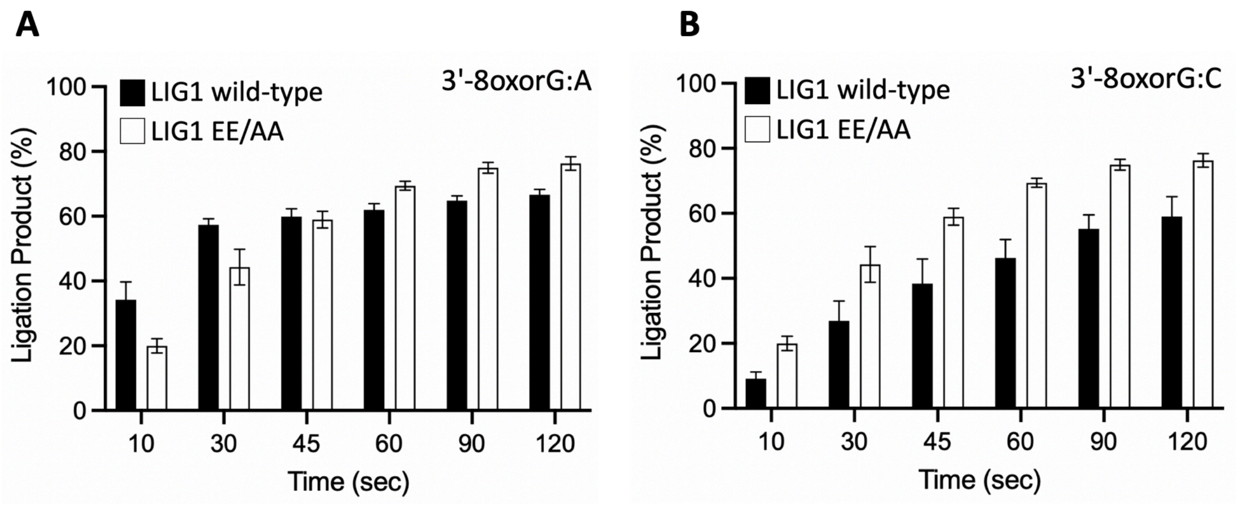


**Supplementary Figure 10. Comparison of the ligation efficiency between LIG1 wild-type and EE/AA mutant.** Graphs show the time-dependent change in the amount of ligation products for nick DNA substrates containing 3'-8oxorG:A (**A**) and 3'-8oxorG:C (**B**) by LIG1 wild-type *versus* EE/AA mutant. The data represent the average from three independent experiments ± SD.

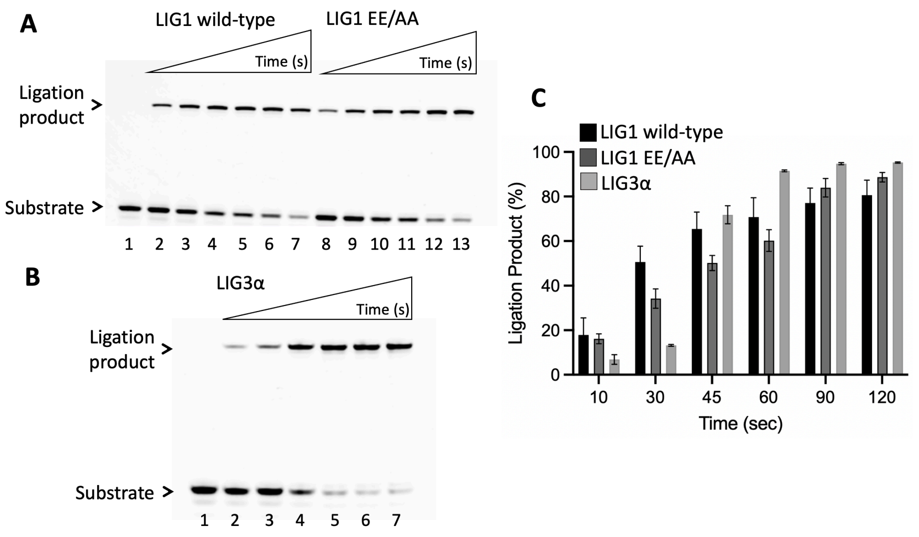


**Supplementary Figure 11. Ligation of the nick DNA substrate with canonical end by LIG1 and LIG3α. (A)** Line 1 is the negative enzyme control of the nick DNA substrate with 3'-dG:C. Lanes 2-7 and 8-13 are the ligation products by LIG1 wild-type and EE/AA mutant, respectively, and correspond to time points of 10, 30, 45, 60, 90, and 120 sec. **(B)** Line 1 is the negative enzyme control of the nick DNA substrate with 3'-dG:C. Lanes 2-7 are the ligation products by LIG3α, and correspond to time points of 10, 30, 45, 60, 90, and 120 sec. **(C)** Graph shows the time-dependent change in the amount of ligation products and the data represent the average from three independent experiments ± SD.

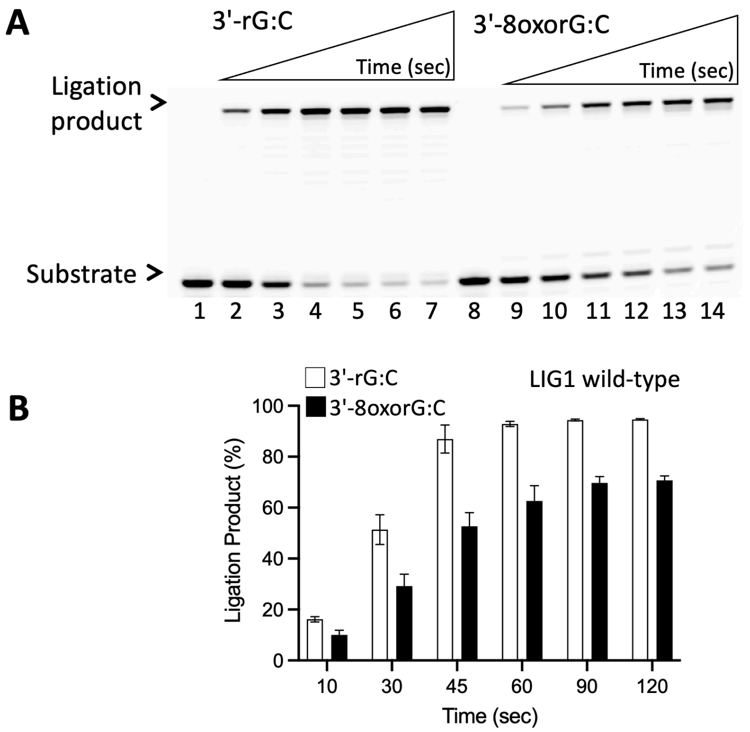


**Supplementary Figure 12. Ligation of the nick DNA substrates with undamaged *versus* damaged ribonucleotide-containing ends by LIG1. (A)** Lanes 1 and 8 are the negative enzyme controls of the nick DNA substrates, and lanes 2-7 and 9-14 are the ligation products by LIG1 in the presence of 3'-rG:C and 3'-8oxorG:C, respectively, and correspond to time points of 10, 30, 45, 60, 90, and 120 sec. **(B)** Graph shows the time-dependent change in the amount of ligation products and the data represent the average from three independent experiments ± SD.

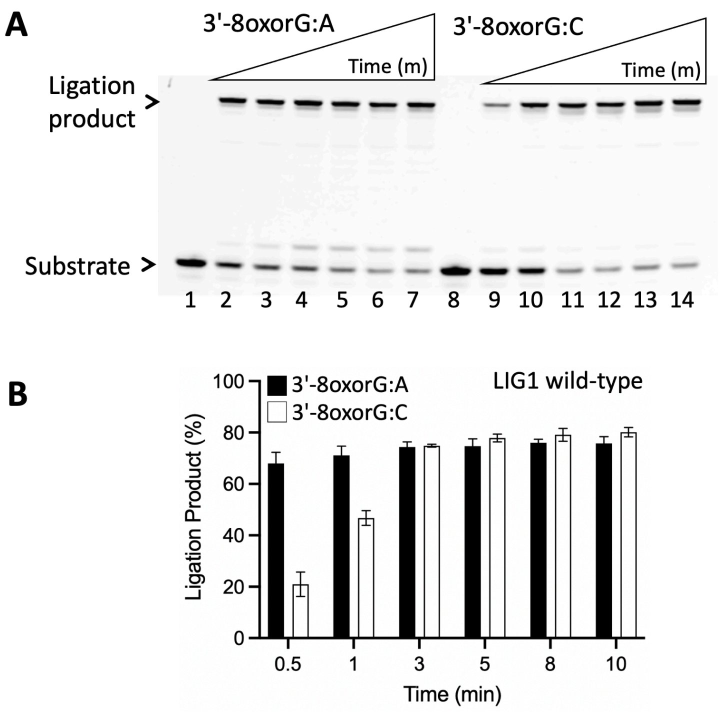


**Supplementary Figure 13. Ligation of the nick DNA substrate with 8oxorG by LIG1. (A)** Lanes 1 and 8 are the negative enzyme controls of the nick DNA substrates with 3'-8oxorG:A and 3'-8oxorG:C, respectively. Lanes 2-7 and 9-14 are the ligation products by LIG1 wild-type in the presence of 3'-8oxorG:A and 3'-8oxorG:C, respectively, and correspond to time points of 0.5, 1, 3, 5, 8, and 10 min. **(B)** Graph shows the time-dependent change in the amount of ligation products and the data represent the average from three independent experiments ± SD.

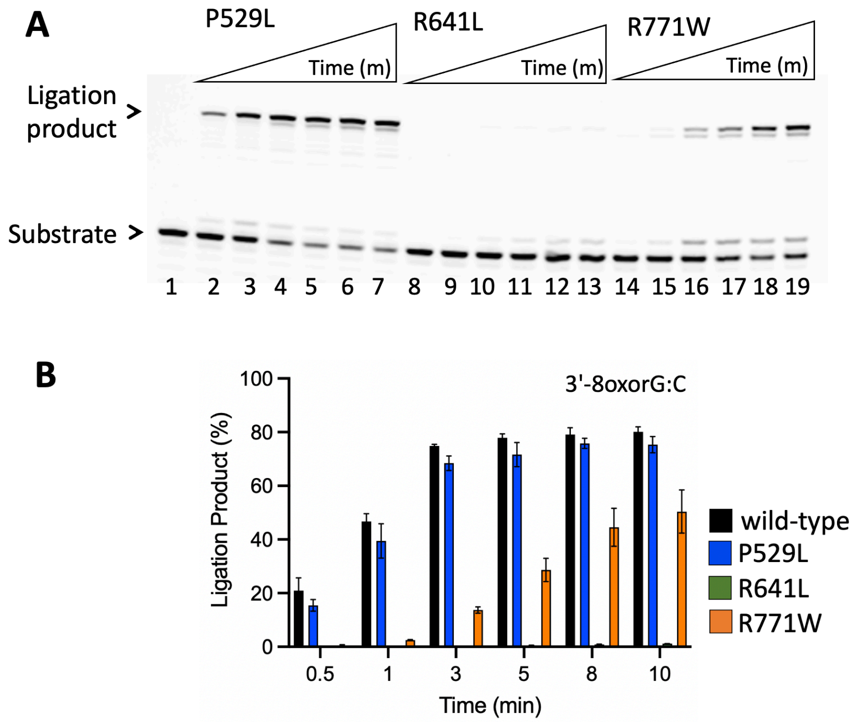


**Supplementary Figure 14. Ligation of the nick DNA substrates with damaged ribonucleotide-containing end by LIG1 deficiency disease mutants. (A)** Lanes 1 is the negative enzyme control of the nick DNA substrate with 3'-8oxorG:C, and lanes 2-7, 8-13, and 14-19 are the ligation products by LIG1 variants P529L, R641L, and R771W, respectively, and correspond to time points of 0.5, 1, 3, 5, 8, and 10 min. **(B)** Graph shows the time-dependent change in the amount of ligation products and the data represent the average from three independent experiments ± SD.

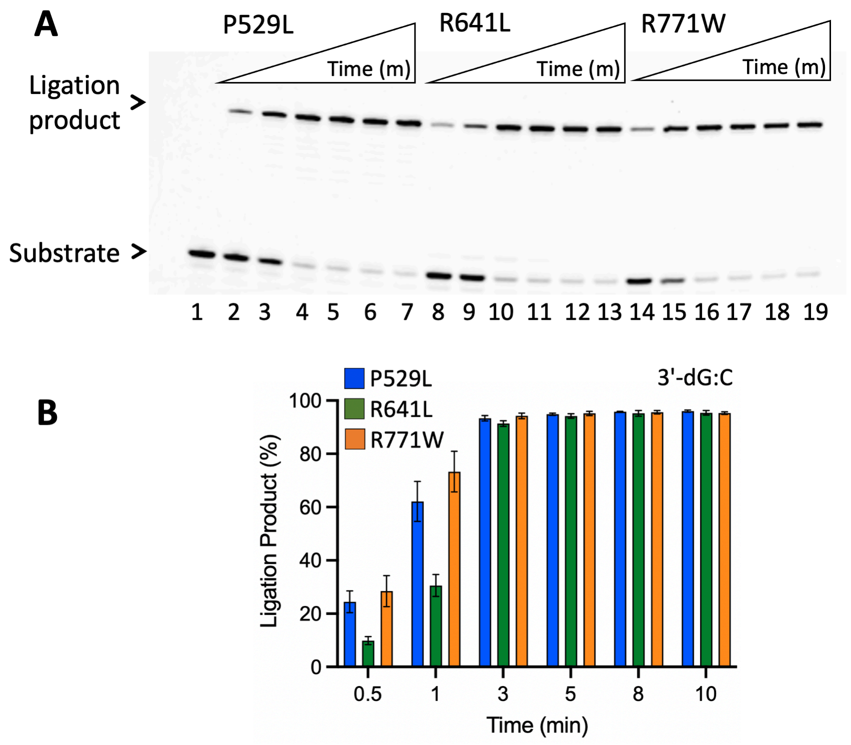


**Supplementary Figure 15. Ligation of the nick DNA substrate with canonical end by LIG1 deficiency disease mutants. (A)** Line 1 is the negative enzyme control of the nick DNA substrate with 3'-dG:C. Lanes 2-7, 8-13, and 14-19 are the ligation products by LIG1 variants P529L, R641L, and R771W, respectively, and correspond to time points of 0.5, 1, 3, 5, 8, and 10 min. **(B)** Graph shows the time-dependent change in the amount of ligation products and the data represent the average from three independent experiments ± SD.

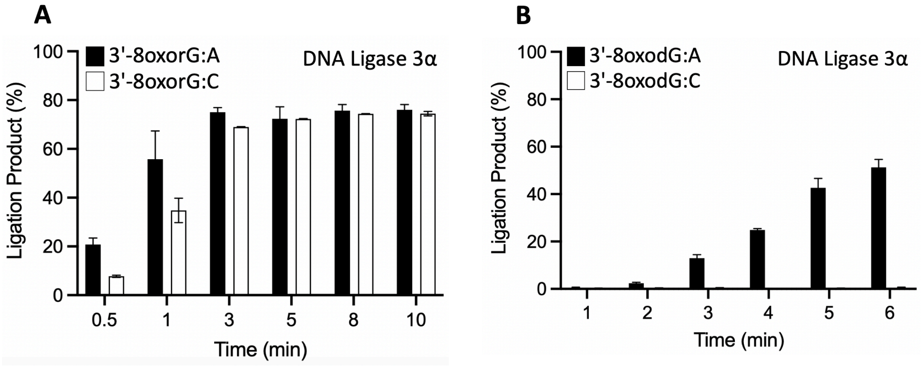


**Supplementary Figure 16. Comparison of the ligation efficiency in the presence of oxidatively damaged ends by LIG3α.** Graphs show the time-dependent change in the amount of ligation products for nick DNA substrates containing 3'-8oxorG:A *versus* 3'-8oxorG:C (**A**) 3'-8oxodG:A *versus* 3'-8oxodG:C (**B**) by LIG3α. The data represent the average from three independent experiments ± SD.

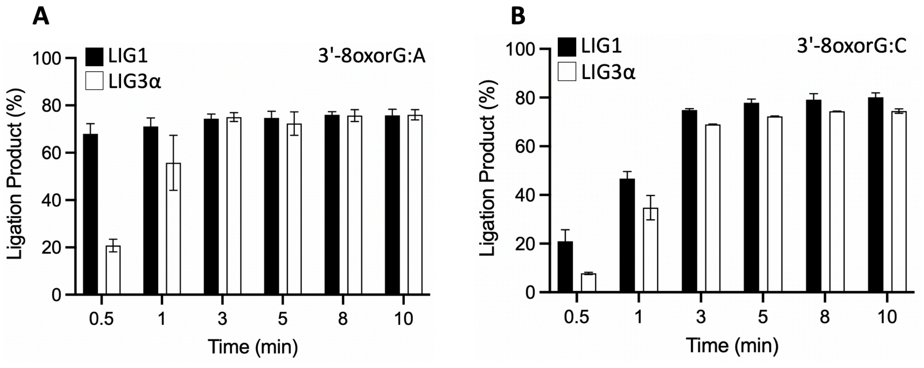


**Supplementary Figure 17. Comparison of the ligation efficiency in the presence of oxidatively damaged ends by LIG1 *versus* LIG3α.** Graphs show the time-dependent change in the amount of ligation products for nick DNA substrates containing 3'-8oxorG:A (**A**) 3'-8oxorG:C (**B**) by LIG1 *versus* LIG3α. The data represent the average from three independent experiments ± SD.

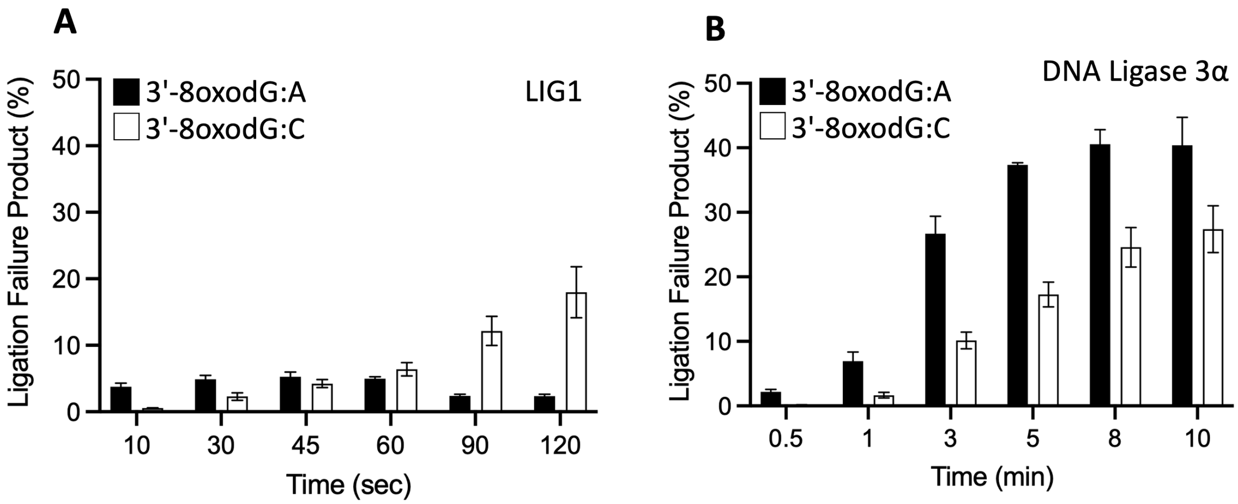


**Supplementary Figure 18. Comparison of the ligation failure in the presence of 3'-8oxodG by LIG1 *versus* LIG3α.** Graphs show the time-dependent change in the amount of ligation failure products (DNA-AMP) in the presence of nick DNA substrates containing 3'-8oxodG:A and 3'-8oxodG:C by LIG1 (**A**) *versus* LIG3α (**B**). The data represent the average from three independent experiments ± SD.

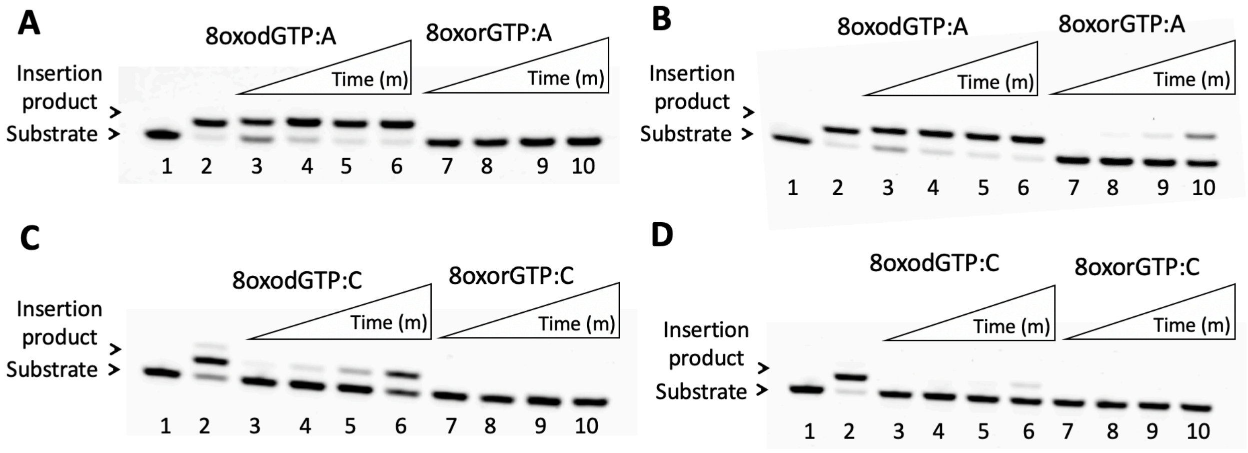


**Supplementary Figure 19. Insertion efficiencies of oxidized nucleotides by polβ. (A-B)** Line 1 is the negative enzyme control of gap DNA substrate with template base A. Lanes 2-6 are the 7-10 are 8oxodGTP:A and 8oxorGTP:A insertion products, respectively, by polβ wild-type (A) and Y271G mutant (B), and correspond to time points of 0.5, 1, 3, 5, 8, and 10 min. **(C-D)** Line 1 is the negative enzyme control of gap DNA substrate with template base C. Lanes 2-6 are the 7-10 are 8oxodGTP:C and 8oxorGTP:C insertion products, respectively, by polβ wild-type (C) and Y271G mutant (D), and correspond to time points of 0.5, 1, 3, 5, 8, and 10 min.

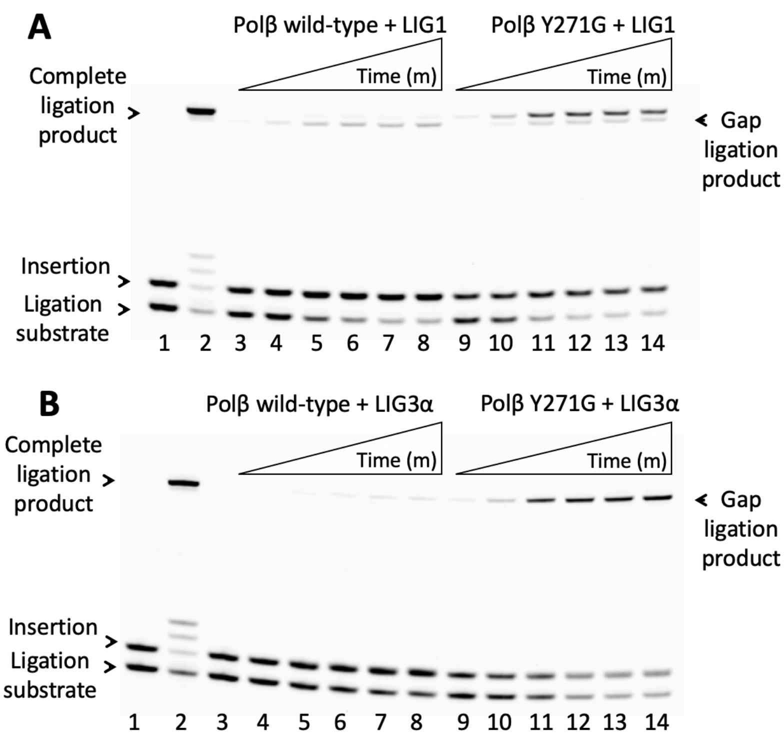


**Supplementary Figure 20. Ligation of polβ 8oxorGTP insertion products by LIG1 and LIG3α. (A-B)** Line 1 is the negative enzyme control of gap DNA substrate with template A. Line 2 is the positive control for the ligation of polβ correct dGTP:C insertion product by LIG1 (A) and LIG3α (B). Lanes 3-8 (polβ wild-type) and 9-14 (polβ Y271G mutant) are the ligation of 8oxorGTP:A insertion products by LIG1 (A) and LIG3α (B), and correspond to time points of 0.5, 1, 3, 5, 8, and 10 min.

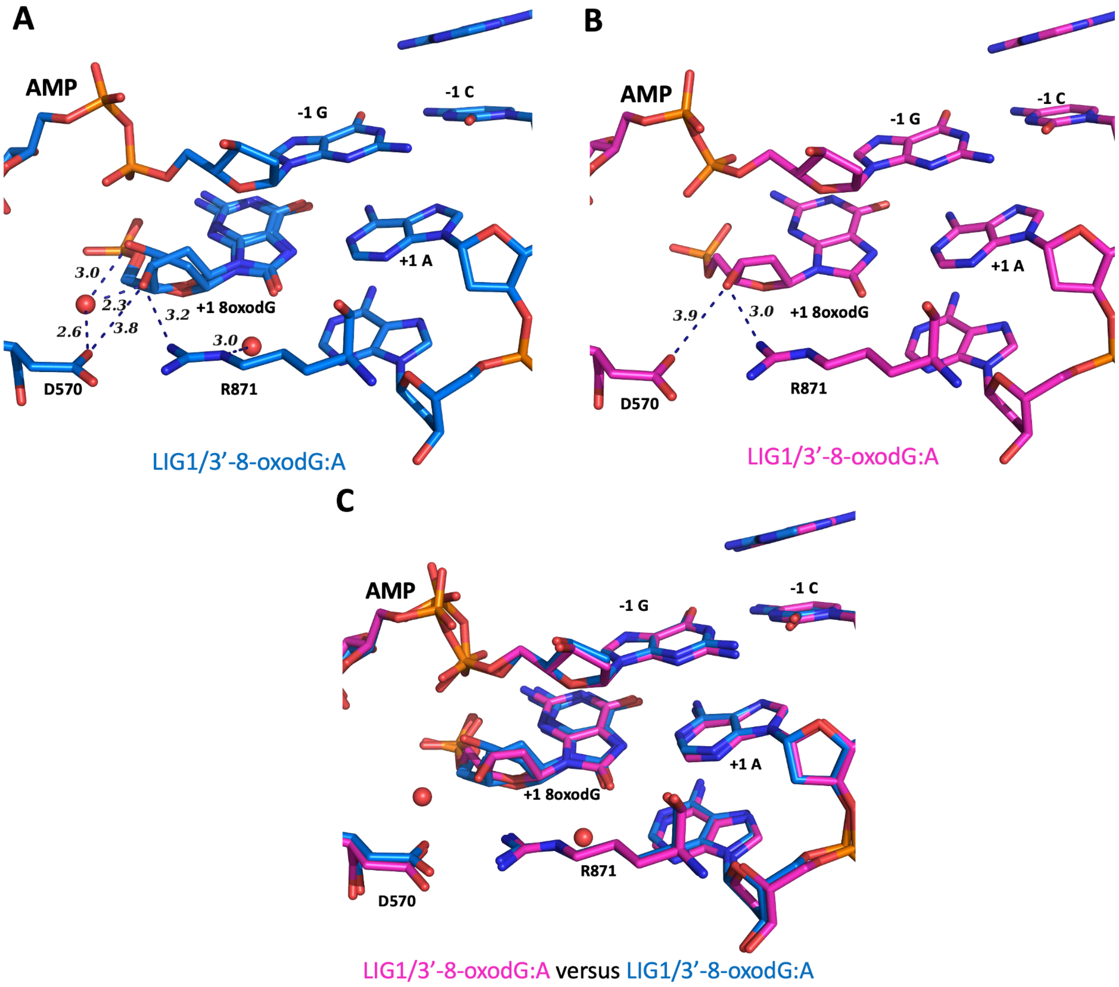


**Supplementary Figure 21. Overlays of LIG1/nick DNA structures containing 8oxodG.** The superimposition of LIG1 structures are shown for 3'-8oxodG:A showing step 2 of the ligation reaction where AMP is bound to 5'-$PO_4$ of nick. Previously solved structure (PDB: 6P0E, 51) by other group (x) is used for this overlay with our LIG1/3'-8oxodG:A structure.

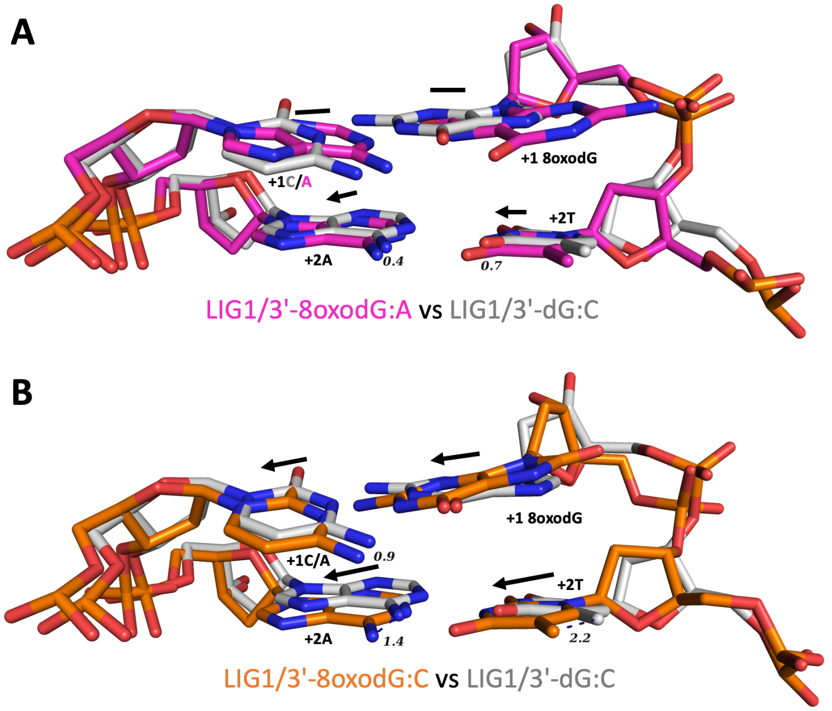


**Supplementary Figure 22. Overlays of LIG1/nick DNA structures with and without damaged ends.** The superimposition of LIG1 structures are shown for 3'-dG:C with 3'-8oxodG:A or 3'-8oxodG:C and demonstrates movements at the +2 nucleotides (+2A and +2T) relative to 3'-OH strand of nick. We previously solved structure of LIG1/3'-dG:C (PDB:8VDN, 53).

| Oligonucleotide | Sequence (5'-3') |
|---|---|
| Template A | GTCCGACAACGCATCAGC |
| Template C | GTCCGACCACGCATCAGC |
| Upstream X<br>(3'-8oxodG or 3'-8oxorG) | GCTGATGCGT**X** |
| Downstream (5'-P) | P-GTCGGAC |

**Supplementary Table 1. Oligonucleotides used in LIG1 crystallization.** Upstream oligonucleotides including a damaged base at the 3'-end (8oxodG or 8oxorG), downstream oligo with phosphate (P) at the 5'-end, and the oligonucleotides containing A or C on template position were used to prepare the nick DNA substrates for LIG1 crystallizations. The base at template base position is underlined and the base position at the 3'-end of nick is shown in bold.

| Nick DNA Substrates | Sequence |
|---|---|
| 3'-dG:C | 5’-CATGGGCGGCATGAACC**G**GAGGCCCATCCTCACC-3’-FAM<br>3’-GTACCCGCCGTACTTGGCCTCCGGGTAGGAGTGG-5’ |
| 3'-8oxodG:A | 5’-CATGGGCGGCATGAACC**X**GAGGCCCATCCTCACC-3’-FAM<br>3’-GTACCCGCCGTACTTGGACTCCGGGTAGGAGTGG-5’ |
| 3'-8oxodG:C | 5’-CATGGGCGGCATGAACC**X**GAGGCCCATCCTCACC-3’-FAM<br>3’-GTACCCGCCGTACTTGGCCTCCGGGTAGGAGTGG-5’ |
| 3'-8oxorG:A | 5’-CATGGGCGGCATGAACC$^{r}$**X**GAGGCCCATCCTCACC-3’-FAM<br>3’-GTACCCGCCGTACTTGGACTCCGGGTAGGAGTGG-5’ |
| 3'-8oxorG:C | 5’-CATGGGCGGCATGAACC$^{r}$**X**GAGGCCCATCCTCACC-3’-FAM<br>3’-GTACCCGCCGTACTTGGCCTCCGGGTAGGAGTGG-5’ |

**Supplementary Table 2. Nick DNA substrates used in ligation assays.** FAM denotes a fluorescence tag and is located at the 3'-end of the nick DNA substrates. The base at 3'-end is shown as bold and the template base is underlined.

| Gap DNA Substrates | Sequence |
|---|---|
| Gap A | FAM-5’-CATGGGCGGCATGAACC GAGGCCCATCCTCACC-3’<br>3’-GTACCCGCCGTACTTGGACTCCGGGTAGGAGTGG-5’ |
| Gap C | FAM-5’-CATGGGCGGCATGAACC GAGGCCCATCCTCACC-3’<br>3’-GTACCCGCCGTACTTGGCCTCCGGGTAGGAGTGG-5’ |

**Supplementary Table 3. Gap DNA substrates used in nucleotide insertion assays.** FAM denotes a fluorescence tag and is located at the 5'-end of the gap DNA substrates. The template base is underlined.

| Gap DNA Substrates | Sequence |
|---|---|
| Gap A | FAM-5’-CATGGGCGGCATGAACC GAGGCCCATCCTCACC-3’-FAM<br>3’-GTACCCGCCGTACTTGGACTCCGGGTAGGAGTGG-5’ |

**Supplementary Table 4. Gap DNA substrate used in coupled assays.** FAM denotes a fluorescence tag and is located at both 3'- and 5'-ends of the gap DNA substrate. The template base is underlined.

| Nick DNA Substrates | Sequence |
|---|---|
| 3'-8oxorG:A | FAM-5’-CATGGGCGGCATGAACC$^{r}$**X**GAGGCCCATCCTCACC-3’<br>3’-GTACCCGCCGTACTTGGACTCCGGGTAGGAGTGG-5’ |
| 3'-8oxorG:A | FAM-5’-CATGGGCGGCATGAACC$^{r}$**X**GAGGCCCATCCTCACC-3’- FAM<br>3’-GTACCCGCCGTACTTGGACTCCGGGTAGGAGTGG-5’ |

**Supplementary Table 5. Nick DNA substrates used in APE1 assays.** FAM denotes a fluorescence tag, the base at 3'-end is shown as bold and the template base is underlined.

| **RMSD (Å)** | LIG1/3'-8oxodG:A Step 2 | LIG1/3'-8oxodG:C Step 2 | LIG1/3'-8oxorG:A Step 3 | LIG1/3'-8oxorG:C Step 2 | LIG1/3'-rG:C Step 3 | LIG1/3'-dG:C Step 2 | LIG1/3'-rG:C Step 2 |
|---|---|---|---|---|---|---|---|
| LIG1/3'-8oxodG:A Step 2 | | 0.237 | 0.513 | 0.313 | 0.437 | 0.436 | 0.605 |
| LIG1/3'-8oxodG:C Step 2 | | | 0.572 | 0.326 | 0.448 | 0.441 | 0.601 |
| LIG1/3'-8oxorG:A Step 3 | | | | 0.628 | 0.602 | 0.544 | 0.472 |
| LIG1/3'-8oxorG:C Step 2 | | | | | 0.477 | 0.511 | 0.654 |
| LIG1/3'-rG:C Step 3 | | | | | | 0.392 | 0.391 |
| LIG1/3'-dG:C Step2 | | | | | | | 0.335 |

**Supplementary Table 6.** The root mean square deviation (RMSD) of LIG1 structures solved previously and presented in this study.

| **DNA ligase 1** | **DNA** | **Ligation Step** | **Reference** |
|---|---|---|---|
| LIG1$^{EE/AA}$ | 3'-dC:G | Step 2 | 51 |
| LIG1$^{EE/AA}$ | 3'-8oxodG:A | Step 2 | 51 |
| LIG1$^{EE/AA}$ | 3'-dA:T | Step 2 | 52 |
| LIG1$^{EE/AA}$ | 3'-dG:T | Step 2 | 52 |
| LIG1$^{EE/AA}$ | 3'-dA:C | Step 1 | 52 |
| LIG1$^{EE/AA}$ | 3'-dG:C | Step 2 | 53 |
| LIG1$^{EE/AA}$ | 3'-rA:T | Pre-step 3 | 53 |
| LIG1$^{EE/AA}$ | 3'-rG:C | Pre-step 3 | 53 |
| LIG1$^{EE/AA}$ | 3'-rA:T | Post-step 3 | 53 |
| LIG1$^{EE/AA}$ | 3'-rG:C | Post-step 3 | 53 |
| LIG1$^{EE/AA}$ | 3'-8oxodG:A | Step 2 | Present study |
| LIG1$^{EE/AA}$ | 3'-8oxodG:C | Step 2 | Present study |
| LIG1$^{EE/AA}$ | 3'-8oxorG:A | Step 3 | Present study |
| LIG1$^{EE/AA}$ | 3'-8oxorG:C | Step 2 | Present study |

**Supplementary Table 7.** The list of LIG1 structures solved previously and presented in this study.

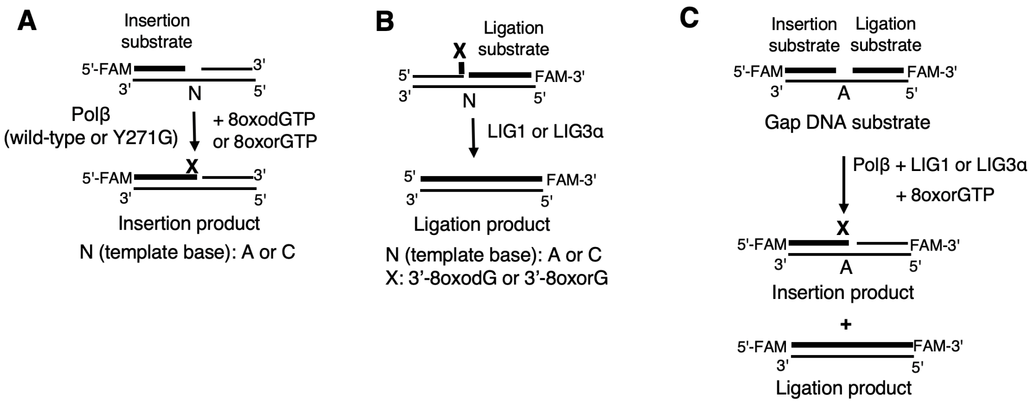


**Supplementary Scheme 1. Illustrations of insertion and ligation assays used in this study**. Schemes show the substrates and reaction products observed in the assays of polβ oxidized nucleotide insertion (A), ligation of nick DNA substrates with oxidatively damaged ends (B), and the coupled assay to measure the ligation of polβ oxidized nucleotide insertion products by DNA ligase simultaneously in the same reaction mixture.

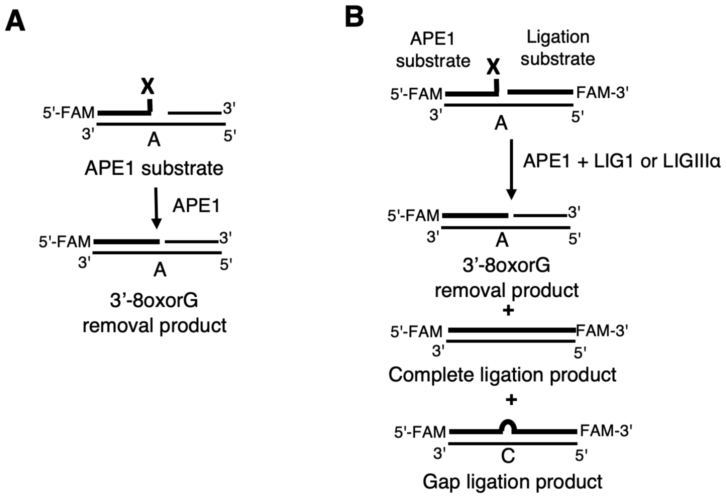


**Supplementary Scheme 2. Illustrations of APE1 assays used in this study**. Schemes show the substrates and reaction products observed in the assays for the removal of oxidized base from the nick DNA substrate by APE1 (A) and the coupled assay to measure the ligation after the removal of oxidized base by APE1 simultaneously in the same reaction mixture (B).
